# ATP-dependent reorientation on a nucleosome primes INO80 for DNA translocation

**DOI:** 10.64898/2026.08.11.744281

**Authors:** Hao Wu, Upneet Kaur, Chengmin Li, Elise N. Muñoz, Geeta J. Narlikar, Yifan Cheng

## Abstract

ATP-dependent chromatin remodeling enzymes play central roles in genome regulation by mobilizing nucleosomes. All remodelers except for one, position their core ATPase at an internal nucleosome location, superhelical location (SHL) 2, and translocate DNA from this site to alter nucleosome conformation. The exception, the remodeler INO80, positions its core ATPase, Ino80, near the DNA exit/entry site, SHL −6. This major architectural difference has raised the question of whether INO80 acts by a fundamentally different mechanism from other remodelers. Here, using cryogenic electron microscopy we capture a new conformation of INO80 that is substantially enriched upon ATP hydrolysis and has Ino80 positioned at an internal nucleosomal location. Comprehensive conformational landscape analysis further uncovers an ATP hydrolysis dependent continuum of additional INO80 conformational states on a nucleosome that were not previously detected. Our studies provide compelling evidence to support a model where INO80 initially engages nucleosomes near SHL −6, followed by a dramatic ATP hydrolysis dependent ~180° reorientation around the nucleosome to place its Ino80 near SHL −2 from where DNA is translocated. INO80’s unique reorientation has broad implications for understanding how ATP-dependent steps that precede nucleosome mobilization can increase the fidelity of remodeling by being responsive to nucleosomal cues.

## Introduction

Eukaryotic DNA is packaged into chromatin, in which approximately 147 bp of DNA wraps around a histone octamer to form a nucleosome, the fundamental repeating unit that organizes the genome and regulates access to DNA^1-3^. ATP-dependent chromatin remodelers play essential roles in regulating chromatin structure for all DNA dependent processes such as transcription, DNA replication, DNA repair and recombination. These machines drive a variety of biochemical outcomes that include nucleosome repositioning, disassembly and histone exchange, and are organized into four major families: SWI/SNF, ISWI, CHD and INO80^4-7^. Structurally, upon nucleosome binding, most remodelers are found to predominantly position their catalytic ATPase subunit at an internal location on nucleosomal DNA, named superhelical location (SHL) −2 or +2 (Figure 1a)^8-13^. This positioning of the ATPase is consistent with biochemical studies indicating that these remodelers translocate DNA from SHL +/−2 to drive nucleosome sliding^14^. In contrast, the INO80 complex from the INO80 family is found to bind a nucleosome with its ATPase subunit positioned near the entry/exit site of the nucleosomal DNA, SHL-6 or SHL+6 (ref 15-18). This difference has suggested that INO80 functions by a distinct mechanism, where its ATPase subunit engages and translocates DNA from SHL −6 (Supplementary Figure 1a). However, recent work has led to an alternative model for INO80 remodeling, wherein ATP hydrolysis drives a large reorientation of INO80, positioning its ATPase motor at SHL −2, from where DNA translocation is catalyzed (Figure 1b)^17^. Distinguishing between these models is fundamental to understanding how INO80’s essential activities are deployed and regulated *in vivo*.

**Figure 1.**
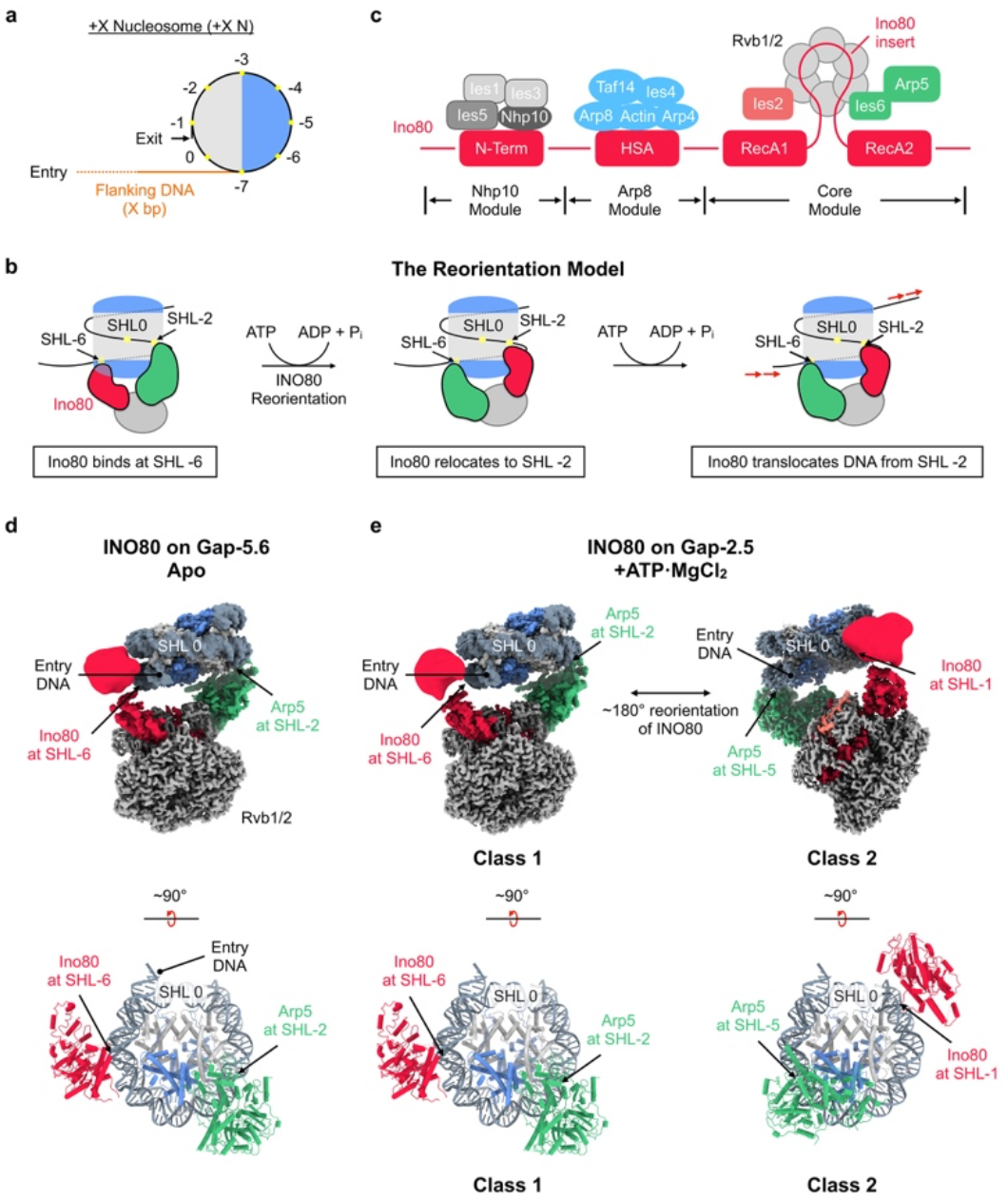
Structures with Gap-2.5 nucleosomes reveal a large reorientation of INO80 upon ATP hydrolysis. Cartoon illustration of a nucleosome with X bp of flanking DNA. The H2A-H2B dimer proximal to the flanking DNA (entry-side dimer) is shown in cornflower blue; H3-H4, light gray; DNA, black; and flanking DNA, orange. DNA Super Helical Locations (SHLs) are marked as yellow dots and the corresponding numbers are in black. **b)** The reorientation model of INO80-mediated nucleosome remodeling. **c)** Architecture of the INO80 complex with different modules shown. **d)** Composite cryo-EM map of INO80/Gap-5.6 complex in apo state (upper panel). The corresponding atomic model is shown below in a view rotated by 90°. H2A-H2B dimers are colored blue; H3-H4, light gray; DNA, light slate gray; Ino80, red; Arp5/Ies6, green; and Rvb1/2, dark gray. The nucleosome dyad (SHL 0) is labeled. **e)** Composite cryo-EM density maps of INO80/Gap-2.5 nucleosome complex in the presence of ATP, revealing two distinct conformations. The corresponding atomic models are shown below in views rotated by 90°. With nucleosome in the same orientation, INO80 in Class 2 is oriented ~180° from INO80 in Class 1. Domains are colored same as in **(d)**. The nucleosome dyad (SHL 0) is labeled.

The alternative model, which we term the reorientation model, was inspired by integrating three types of data with *Saccharomyces cerevisiae* (*S. c*.) INO80. The first type of data involves hexasomes, which are subnucleosomal particles missing an H2A-H2B dimer and proposed to form when RNA polymerase transcribes through a nucleosome. Hexasomes are also active substrates for INO80 (ref 19). On hexasomes, unlike on nucleosomes, the ATPase subunit of INO80 is positioned near SHL −2 (ref 17,20). The second type of data involves a classical biochemical assay used to identify the DNA site from where a remodeler translocates nucleosomal DNA^21-23^. In this assay a single nucleotide gap is placed at defined locations in nucleosomal DNA and the effects on remodeling are assessed. If a gap at a given location inhibits remodeling, this is interpreted as ATPase action at that location being important for DNA translocation. In such studies it was found that placing a gap near SHL −6 (SHL −5.6) and near SHL −2 (SHL −2.5) both reduce INO80 remodeling activity on nucleosomes by ~200-fold^17,20,24^. The third type of data involves single molecule FRET experiments monitoring the sliding of nucleosomes. These experiments identified an ATP-dependent pause that occurs before ATP-dependent DNA translocation, implying two different effects of ATP hydrolysis^25^. This pause could reflect that a reorientation of INO80 is a necessary ATP-dependent process prior to ATP-dependent DNA translocation. Together these data are consistent with the possibility that, on nucleosomes, the ATPase subunit of INO80 needs to access the SHL −2 location for effective DNA translocation.

However, the biochemical data using DNA gaps does not definitively distinguish between the two models. This is because nucleosomal DNA gaps could also affect the action of non-ATPase subunits within the multi-subunit INO80 complex. This is exemplified by *S. c*. INO80, a multi-module complex that is assembled on the Ino80 subunit, which has an N-terminal region, a helicase-SANT-associated (HSA) domain, and a catalytic motor composed of two RecA-like lobes connected by a long insertion. The different regions of the Ino80 protein organize the Nhp10, Arp8, and Core modules (Figure 1c). In the Core module, two RecA lobes come together forming an ATPase domain, Ino80^ATPase^. The hexameric Rvb1/2 ring is assembled on the insertion loop between the two RecA lobes. The Arp5 module is attached to the Rvb1/2 ring on the opposite side of Ino80^ATPase^. In available structures of INO80 bound to a nucleosome, Ino80^ATPase^ is positioned near SHL −6 and the Arp5 module near SHL −2 (ref 15,16,18). Thus, inhibition of remodeling by a gap near SHL −2 could reflect either disruption of ATPase-dependent DNA translocation from SHL−2, as proposed by the reorientation model, or disruption of productive Arp5 engagement at the same location. It has been difficult to resolve this ambiguity using current biochemical and structural approaches. Further, direct structural evidence for a reorientation model is lacking.

In this study, using single particle cryogenic electron microscopy (cryo-EM), we compared the conformational landscape of how INO80 engages a nucleosome in the absence versus presence of ATP using substrates with and without single nucleotide gaps. In the presence of ATP, we observe an ensemble of new conformations of INO80 bound around the nucleosome, many of which include Ino80^ATPase^ positioned near SHL −2. These data provide direct structural support for the reorientation model. Our findings further explain how ATP hydrolysis is used in two different ways by INO80: to reorient Ino80^ATPase^ from SHL −6 towards SHL −2 and to translocate DNA from SHL −2. The use of ATP for the reorientation of INO80 indicates that regulating this step is of high biological significance and likely makes INO80 responsive to nucleosomal features near SHL −6. More broadly, our work highlights how active conformations of a complex machine such as INO80 may only be sufficiently populated during ATP-hydrolysis and difficult to capture by conventional ATP analog based methods.

## Results

### INO80 undergoes a dramatic reorientation on a nucleosome upon ATP hydrolysis

We used nucleosomes with DNA gaps near SHL-6 and SHL-2 to structurally test the reorientation model. If ATP hydrolysis drives reorientation of Ino80^ATPase^ to SHL −2, a prediction is that a DNA gap placed between SHL −6 and −2 would be translocated up to SHL −2, accumulating at that location because it would inhibit further DNA translocation by Ino80^ATPase^. Correspondingly, placing a gap near SHL −2 may trap the reoriented Ino80^ATPase^ bound near SHL −2, by inhibiting translocation. In contrast, if Ino80^ATPase^ translocates DNA from SHL −6, then upon adding ATP, Ino80^ATPase^ is predicted to remain bound near SHL −6 regardless of whether the DNA gap is at SHL −2, SHL −6 or in between.

To test these predictions, we reconstituted nucleosomes using the 601 positioning sequence with 80 bp of flanking DNA and introduced a single nucleotide gap at three different SHLs: −2.5, −3.6 and −5.6, hereafter referred to as Gap-2.5, Gap-3.6 and Gap-5.6, respectively (Supplementary Figure 1b)^26^. We found that INO80 hydrolyzes ATP at similar rates on nucleosomal substrates with or without gaps, indicating that these substrates are competent to fully stimulate INO80’s ATPase activity (Supplementary Figure 1c). Nucleosomes with gaps were incubated with *S. c*. INO80 and ATP for 30 minutes before freezing the cryo-EM grids (Supplementary Figure 1d). For nucleotide free controls (apo), we also determined structures of INO80 bound to Gap-3.6 and −5.6 nucleosomes in the absence of ATP. We collected single particle cryo-EM dataset for each sample individually (Supplementary Figure 1e), merged all five datasets and processed them together as a single one (Supplementary Figure 2). Such processing prevents any potential bias in classification and multi-body refinement of individual datasets separately. After this procedure, particles of each individual dataset can be extracted for further analysis. As reported in our previous studies, classification was followed by focused refinement of the INO80 core and nucleosome separately. This multi-body refinement approach allowed us to generate composite maps of the INO80 core-nucleosome complex and assign the position of Ino80^ATPase^ on the nucleosome (Supplementary Figures 2 and 3).

Consistent with previous studies on non-gapped (also called WT) nucleosomes without ATP^17^, there is only one dominant conformation of the INO80-nucleosome complex from the apo Gap-3.6 and Gap-5.6 datasets, in which the Ino80^ATPase^ binds near SHL −6 while the Arp5 module engages nucleosomes near SHL −2 (Figure 1d). In contrast, when ATP is added to a sample of INO80 with Gap-2.5, two distinct conformations were identified (Figure 1e and Supplementary Figures 2 and 4): One shows Ino80^ATPase^ bound near SHL −6 (Class 1), which is very similar to the structures found from the apo datasets. The other, Class 2, shows Ino80^ATPase^ bound near SHL −1 and the Arp5 module bound near SHL −5, which is almost opposite in orientation relative to INO80-nucleosome complexes in Class 1. Notably, this conformation has not been observed in any previously reported apo or ADP/BeF_x_ bound INO80-nucleosome structures. Further, for Gap-3.6 and Gap-5.6, addition of ATP also resulted in the same two major classes. Together, these findings indicate that Ino80^ATPase^ does not remain steadily bound near SHL −6 during ATP hydrolysis. Instead, the INO80 complex does reorient on the nucleosome in an ATP-dependent manner to bring Ino80^ATPase^ near SHL −2.

### Landscape of relative orientation between INO80 and the nucleosome

The results above raised the question of how INO80 reorients from SHL −6 to SHL −1. Classification and multi-body refinement group the conformationally heterogeneous INO80-nucleosome particles into discrete conformational subgroups. Any less populated intermediates may therefore be averaged out during conventional classification approaches. Indeed, the need for applying multi-body refinement to achieve high resolution individually for INO80 and the nucleosome suggests considerable underlying conformational variations between INO80 and the nucleosome. Additionally, identification of a large Class 2 population in the samples with ATP raised the question of whether a Class 2 like conformation also exists in the absence of ATP but is much less populated. To better understand how INO80 reorients around the nucleosome and to capture the population distributions before and after adding ATP, we applied cryoROLE^27^ to analyze the landscape of relative orientations between the main body of INO80 and the bound nucleosome, assuming both behave as “rigid bodies” during this reorientation process. This method was recently developed and validated in analyzing the landscape of the INO80-hexasome complex where we had captured three distinct substates by conventional multi-body refinement and classification. These substates had Ino80^ATPase^ engaged near, SHL-3, SHL-2.5 and SHL-2 (ref 17). Instead of discrete conformational substates, the landscape analysis revealed a more continuous distribution of INO80 on the hexasome wherein Ino80^ATPase^ occupies multiple positions between SHL −3 to −1 with the predominant location near SHL −2 (ref 27).

Here, we analogously applied cryoROLE to extract the conformational continuum underlying INO80’s engagement with a nucleosome that gets buried during conventional classification (Figure 2 and Supplementary Figure 5). For comparison, we also included the dataset from our previous studies^17,18^ of INO80 bound to a nucleosome without a gap, without added nucleotide or with ADP/BeF_x_, into the landscape analysis (Figure 2 and Supplementary Figure 6). The landscape of relative orientations between INO80 and the nucleosome is depicted in a Cartesian coordinate system, in which α is designated as the angle around the axis perpendicular to the disk-like face of the nucleosome and 0° corresponds to Ino80^ATPase^ at SHL 0, or the dyad. The β angle is around an axis lying in the disk-like face in the direction from where Ino80^ATPase^ contacts DNA to where the Arp5 module contacts DNA. The remaining axis is then defined as the γ angle (Figure 2a and Supplementary Figure 5a). Thus, α describes rotational motion of INO80 around nucleosome, while β and γ describe wobbling of INO80 on the nucleosome at any specific orientation between INO80 and the nucleosome. When particles of all datasets are combined, the landscape is widely spread and distributed continuously along α, with three high particle population hotspots located at SHL −6, −7 and −1 (Supplementary Figure 5b and c, Movie S1). Reconstructions calculated using particles extracted from 15 equally spaced locations in the landscape, with an interval of 22.5° along α from −180° (corresponding to SHL −4) to 100° (SHL −2.2), all show clearly resolved density for the INO80 body along with the Arp5 module and the Ino80^ATPase^, as well as the nucleosome, allowing the Ino80^ATPase^ to be placed on to a specific position on the nucleosome (Supplementary Figure 5d, Movie S2). From this combined dataset, the landscape of each sample can be extracted and compared, which we describe in detail below.

**Figure 2.**
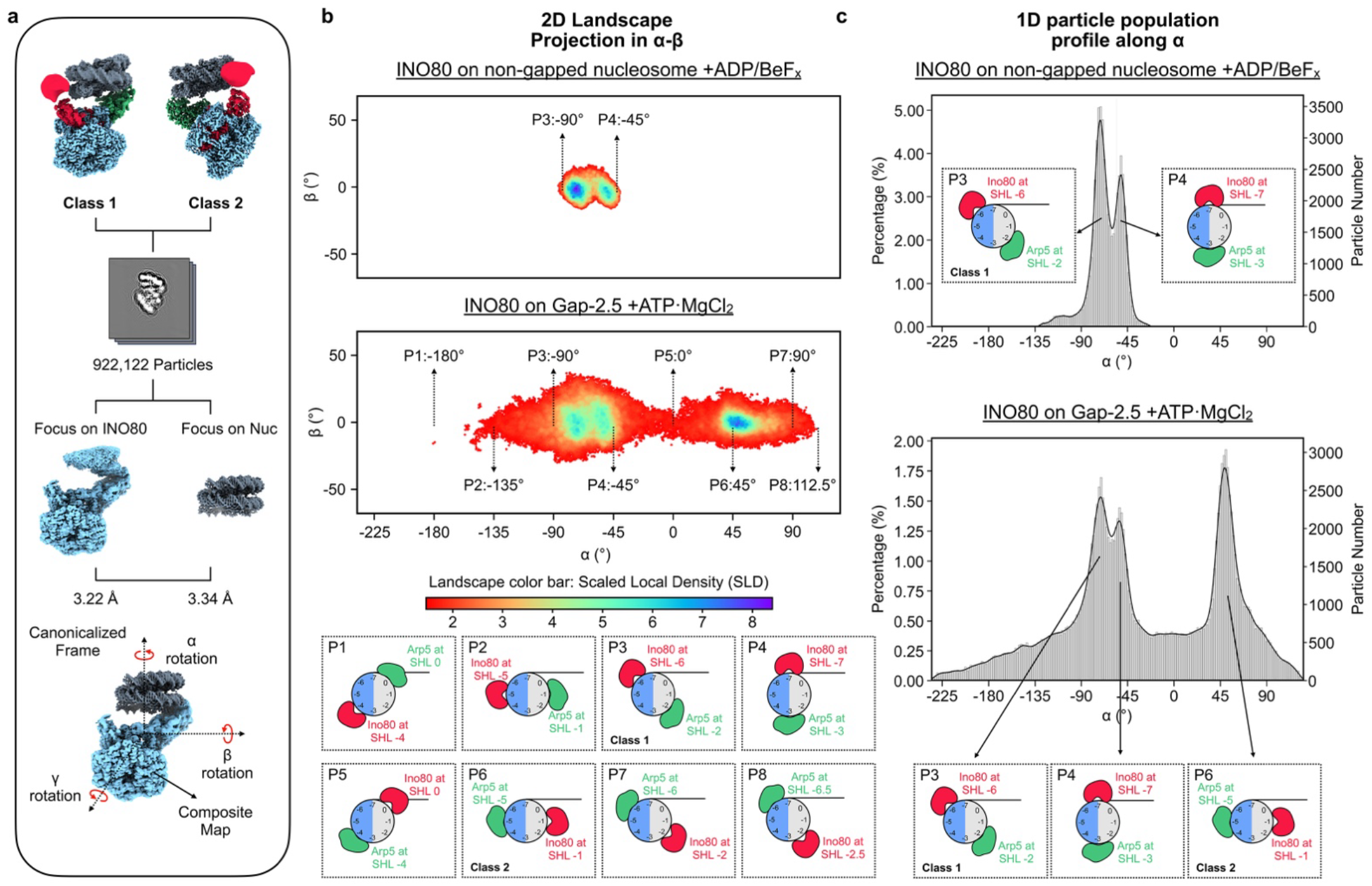
Landscape of the relative orientation between INO80 and the nucleosome. **a)** Schematic illustration of generating a single composite map of the INO80-nucleosome complex. Particles from the two classes were merged and subjected to focused refinements on INO80 and the nucleosome. A composite map is generated by placing Ino80^ATPase^ at SHL 0. Orientation between INO80 and nucleosome is defined by α, β, and γ angles around three perpendicular axes as defined. **b)** Projections of the 3D landscapes onto the α-β plane. Landscape of INO80 bound with non-gapped nucleosome with ADP/BeF_x_ (upper panel) and INO80/Gap-2.5 with ATP (middle panel). Particle distributions are color coded by color bar as scaled local density (SLD), with blue indicating highly populated orientations and red indicating less populated orientations. Bottom panel is cartoon illustration of representative orientations between INO80 and nucleosome selected from the landscape at 45° intervals along the α axis from −180° to 112.5°. Position of Ino80^ATPase^ (red) and Arp5 (green) are marked. A SLD threshold of 1.5 was applied for visualization. **c)** Comparison of particle distribution profile along α derived from the landscapes filtered using an SLD threshold of 1.0. The horizontal axis is α angle, the left y axis indicates the percentage of particles assigned to each α angle relative to the total number of particles (after removing all particles whose SLD is less than 1). The right y axis indicates the corresponding particle number.

### INO80 appears to rotate around the nucleosome upon ATP hydrolysis

The landscapes from all four samples, apo Gap-3.6, apo Gap-5.6, WT nucleosome without nucleotide and WT nucleosome with ADP/BeFx, are similar, with each showing a very confined particle distribution along β and γ, and a narrow spread along α with two hot spots, correlated with the Ino80^ATPase^ located at SHL −6 and −7 (Figure 2b). The two hotspots are consistent with the classification results from previous studies by us and others^15-18^. These results suggest that, without ATP hydrolysis, INO80 largely binds the nucleosome within a narrow range of orientations around axis of α, placing Ino80^ATPase^ preferentially at SHL −6 and −7 (Figure 2b and c). The results also indicate that binding of ADP/BeF_x_ does not substantially change the overall architecture of how INO80 engages with a nucleosome compared to the state where no nucleotide is bound to Ino80^ATPase^. We do observe a small population of particles (~ 1.6% from apo Gap-3.6 and ~ 6% from apo Gap-5.6) that adopt a conformation with Ino80^ATPase^ located near SHL −1, suggesting that this state is much less accessible without ATP (Extended Data Fig 8).

In the presence of ATP, the landscapes from the three gap containing nucleosome samples (Gap-2.5, −3.6 and −5.6) have similar shapes to each other but with some differences in particle distribution profiles (Figure 2b and c, and Supplementary Figure 9). However, they are all strikingly different from the landscapes for the apo and ADP/BeFx containing samples (Supplementary Figures 5 - 9). Taking the landscape of Gap-2.5 with ATP as an example (Figure 2b and c), the particle distribution spans continuously along α from ~ −180° to ~ 112°, with hotspots where Ino80^ATPase^ is located at SHL −6, −7, and −1.

Surprisingly, there are very few particles with Ino80^ATPase^ located between SHL −2.5 to −4. On the opposite side of nucleosome, although at a relatively lower population than in the hotspots, the Ino80^ATPase^ is found near the dyad (Figure 2b and c, and Supplementary Figure 5). In all reconstructions calculated from particles extracted from the landscape (Supplementary Figure 5d), the Arp5 module is well resolved and remains in contact with the nucleosomal DNA upper gyre. In contrast, the density of Ino80^ATPase^ is less well resolved suggesting that it engages with the nucleosomal DNA more dynamically than Arp5 (Supplementary Figure 5c). The reconstructions of Ino80^ATPase^ at SHL −1 and −6/−7 show additional density in between INO80 and the surface of the nucleosome. This density arises from the interaction between a region of Ies6 and the H2B-αC helix (Supplementary Figure 5g). However, this interaction is in the opposite orientation when the Ino80^ATPase^ is bound at SHL −1 compared to SHL −6/−Together, these observations suggest that ATP hydrolysis drives INO80 to rotate around the nucleosome directionally such that, from SHL −6/−7, Ino80^ATPase^ preferentially moves along nucleosomal DNA via the dyad to reach SHL −1, but not via SHL −4 (Supplementary Figure 5e and f). We speculate that the directionality may arise because during INO80’s rotation around the nucleosome, Arp5 needs to consistently maintain contact with the DNA upper gyre. When the Ino80^ATPase^ is at SHL −7, it could potentially swing over the flanking DNA from SHL −7 to SHL −1 while the Arp5 module maintains contact with the DNA upper gyre (Supplementary Figure 5e and f). Rotation of Ino80^ATPase^ in the opposite direction may be unfavorable as such movement would require Arp5 to transiently disengage from DNA to switch from the dyad to SHL −7. Movement of Arp5 towards SHL +1 is also expected to be less favored because of steric hindrance caused by flanking DNA (Supplementary Figure 5e and f).

### The gap is translocated to SHL −2

The new predominant location of INO80 identified with Ino80^ATPase^ near SHL −1 does not readily align with expectations from the reorientation model that Ino80^ATPase^ translocates DNA from SHL −2. To understand where Ino80^ATPase^ translocates DNA from, we asked where the gap in the Gap-3.6 nucleosomes moves to after ATP-dependent remodeling. Structurally, the position of the gap after ATP-dependent remodeling should reveal the position from which DNA is translocated by Ino80^ATPase^. We performed focused refinement and classification on the remodeled nucleosomes using only particles belonging to the Gap-3.6 dataset extracted from Class 2, in which Ino80^ATPase^ engages nucleosomes at SHL −1. A gap can be clearly identified from the refined nucleosome structures and is positioned precisely at SHL −2 (Figure 3a). Consistent with movement of the gap from SHL −3.6 to SHL −2, we observe at least 5 bp of extra DNA on the exit side of the nucleosome (Figure 3b). These findings are consistent with a model where DNA is translocated from SHL −2 by Ino80^ATPase^.

**Figure 3.**
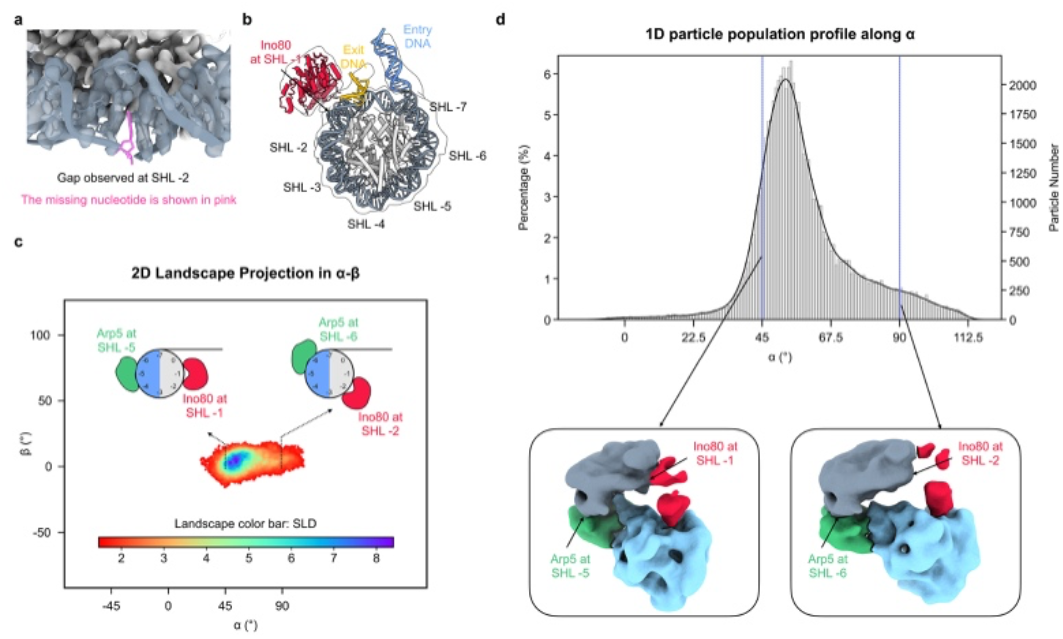
Gap is translocated to SHL −2 upon ATP hydrolysis. Cryo-EM density map of nucleosome overlaid with the atomic model shows a clearly resolved gap at SHL −2, captured from dataset of INO80 with Gap-3.6 and ATP. The missing nucleotide is shown in pink. **b)** Cryo-EM nucleosomal density map is shown as transparent low-resolution contour with the corresponding atomic model docked. Extrusion of the exit DNA is seen. Ino80^ATPase^ is colored red, exit DNA orange, entry DNA blue, histones light gray, and nucleosomal DNA light slate gray. SHLs are labeled. **c)** Landscape shows the distribution of how Ino80^ATPase^ engages nucleosome when gap is trapped at SHL −2. Representative orientations are illustrated as cartoons showing the corresponding Ino80^ATPase^ and Arp5 binding positions on the nucleosome. **d)** Particle distribution profile around axis of α angle derived from the landscape shown in **(c)**. Reconstructions shown below the profile were calculated from particles extracted from the landscape at SHL −1 and −2.

For all particles with the gap at SHL −2, the landscape analysis shows that the position of Ino80^ATPase^ is distributed between SHL −1 and −2, with a peak located closer to SHL −1, and a motion of ~ 30° along β and γ angles, assuming the entire INO80 including Ino80^ATPase^ is one rigid body (Figure 3c and d). Together with the observation that the RecA2 lobe is more poorly resolved than the RecA1 lobe, these variations raise the possibility that the Ino80^ATPase^ has sufficient flexibility to reach the DNA at SHL-2 during active DNA translocation.

### The H2A-H2B acidic patch mutation (APM) reduces INO80 rotation around the nucleosome

The observed ATP-dependent reorientation of INO80 raised the question of how the complex uses its contacts with the nucleosome to facilitate this rotation. Previous structures of INO80 in the apo and ADP-BeF_x_ state, where Ino80^ATPase^ is engaged near SHL −6, have shown that the grappler region of Arp5 makes contacts with the H2A-H2B acidic patch^16,17^. It has also been shown previously that mutation of the H2A-H2B acidic patch (APM) reduces INO80 remodeling activity by ~ 200-fold without affecting ATPase activity, indicating that the acidic patch plays a large role in coupling ATP hydrolysis to nucleosome sliding^16,19^. Interestingly, in both conformations, when Ino80^ATPase^ is near SHL-1 and when it is near SHL −6/−7, regions of INO80 such as Arp5 and Ies6 appear to interact with parts of the acidic patch (Supplementary Figure 5g). To investigate if the acidic patch plays a role in the reorientation of INO80, we performed single particle cryo-EM analysis of INO80 in complex with APM nucleosomes without any DNA gap and with the addition of ATP (Supplementary Figure 10). Like our observations for ATP-dependent conformations of INO80 bound to nucleosomes with DNA gaps, two classes were identified. In Class 1, which contains 90% of particles, Ino80^ATPase^ is found at SHL −6. In Class 2, with only ~10% of particles, Ino80^ATPase^ is found at SHL −1 (Figure 4). The lower population of Class 2 particles with the APM nucleosomes indicates that the acidic patch plays a major role in enabling the reorientation of INO80. Together with the previous finding that APM substantially slows remodeling, these observations further support the model that INO80 translocates DNA from its reoriented conformation. The INO80-acidic patch interaction may promote remodeling in three mutually compatible ways: (i) by stabilizing the interactions in the Class 1 conformation to effectively couple ATP hydrolysis to reorientation, (ii) by lowering the activation barrier for reorientation to enable faster transition between Class 1 and Class 2 conformations and (iii) by stabilizing the Class 2 conformation.

**Figure 4.**
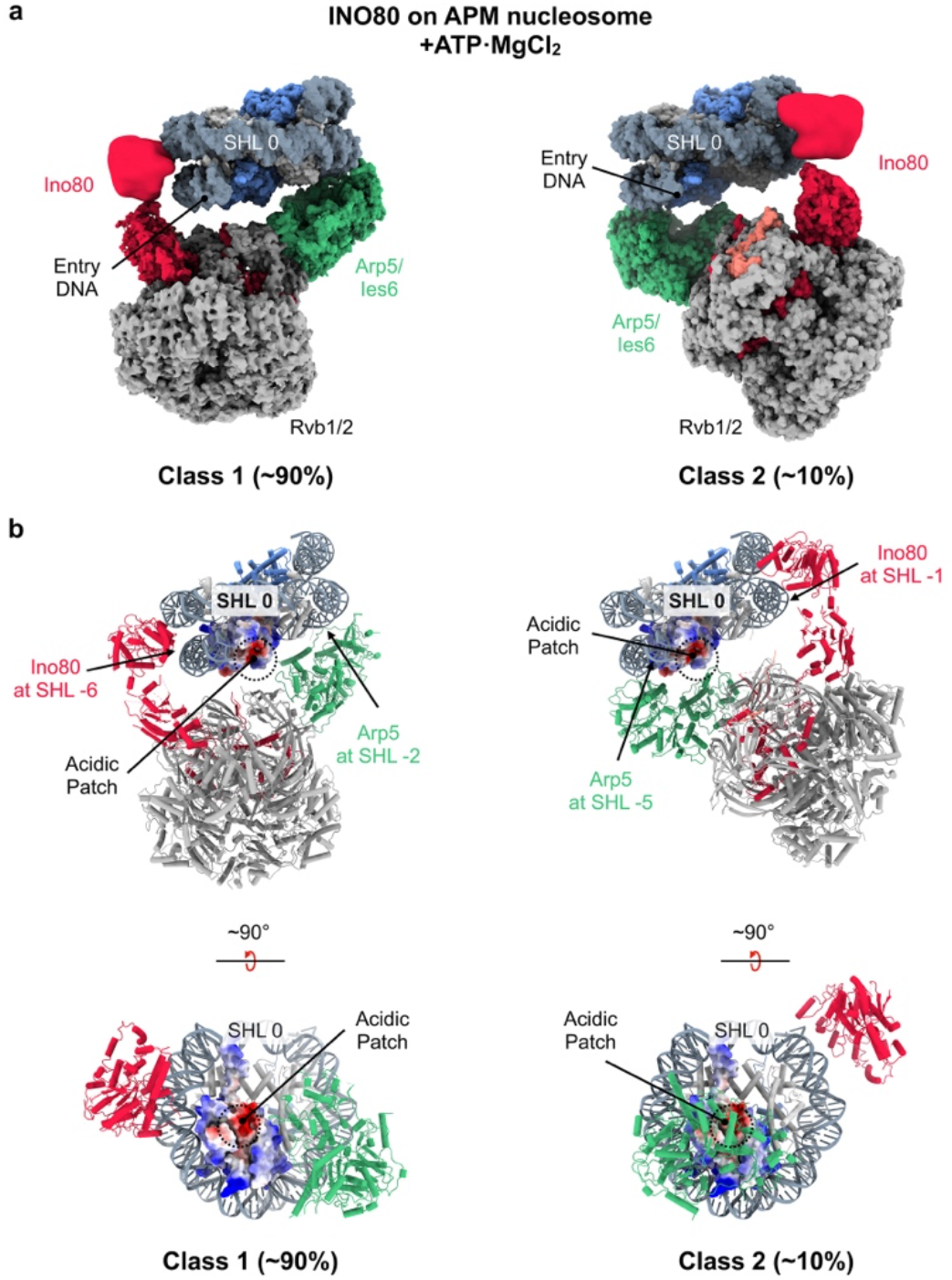
Mutating the acidic patch reduces reorientation of INO80 upon ATP hydrolysis. Cryo-EM density maps of the two classes of INO80/APM nucleosome complex in the presence of ATP, corresponding to Ino80^ATPase^ near SHL −6 (Class 1) and SHL −1 (Class 2). Population of particles in each class are shown as ~90% in Class 1 and ~10% in Class 2. **b)** Corresponding atomic models of the two classes. The upper panels are shown in the same orientations as the cryo-EM maps in **(a)**, and the lower panels are shown in a view rotated by 90°, looking into the nucleosome surface from the orientation of bound INO80. The electrostatic surface representation of the entry-side H2A-H2B dimer is shown, with the acidic patch outlined by the dashed circle.

### A model of how ATP hydrolysis drives both INO80 rotation and DNA translocation

Our data implies that ATP hydrolysis is required for INO80’s rotation around the nucleosome. This raises the question of how Ino80 uses ATP to accomplish two different outcomes: INO80 rotation and DNA translocation. While the landscape analysis suggests that this rotation could be a continuous process, two orientations between INO80 and nucleosome, with Ino80^ATPase^ at SHL −6/−7 and at SHL −1, are more highly populated than those with Ino80^ATPase^ between them. The highly populated states may arise from interactions between INO80, including the Arp5 module and Ies6, with the acidic patch of the nucleosome in addition to interactions made by Arp5 with the nucleosomal DNA. When Ino80^ATPase^ is at SHL −6 and SHL −1, the Arp5 subunit is at SHL −2 and SHL −5 respectively and therefore likely makes different interactions with the acidic patch (Figure 4). Structurally, INO80 shows more conformational variability around the nucleosome when Arp5 is at SHL −2/−3 than when it is at SHL −5, consistent with the possibility that Arp5/Ies6 engages the nucleosome more stably at SHL −5 (Figure 5a and Supplementary Figure 11). We therefore propose that when Ino80^ATPase^ is at SHL −6, the interaction of Arp5/Ies6 with the acidic patch is more susceptible to being dislodged by the Ino80 motor than when the Ino80^ATPase^ is at SHL −1 or SHL −2. As a result, Ino80^ATPase^ action near SHL −6 results in translocation of Ino80 on DNA whereas Ino80^ATPase^ action near SHL −2 results in retention of Ino80^ATPase^ near SHL −2 and translocation of DNA relative to the histone octamer (Figure 5b).

**Figure 5.**
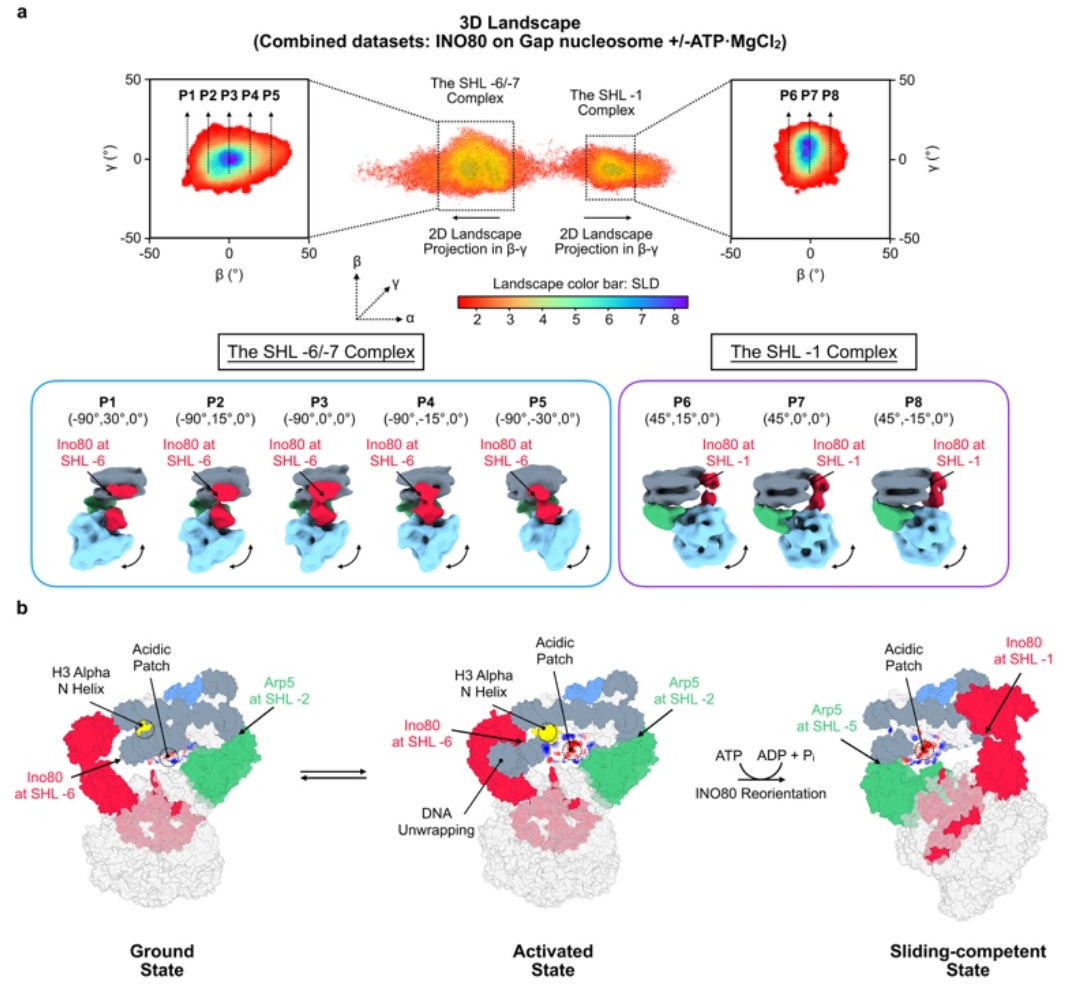
Larger conformational variation of INO80 on nucleosome before reorientation. INO80 shows more conformational variation on a nucleosome when Ino80^ATPase^ is at SHL −6/−7 (left) than when it is near SHL −1, as revealed from the landscape projections (left: SHL −6/−7, right: SHL −1), as well as from reconstructions calculated from particles sampled at 15° intervals along the β axis from the landscape projected onto β-γ. The corresponding reconstructions are shown in the lower panel. **b)** Proposed model for INO80-mediated nucleosome sliding. INO80 initially binds the nucleosome with Ino80^ATPase^ at SHL −6 (ground state). DNA unwrapping activates INO80, and ATP hydrolysis drives INO80 reorientation to reposition Ino80^ATPase^ from SHL −6 to SHL −1, reaching a sliding-competent state that translocates DNA from SHL −2.

Summarizing all data presented in this study, we propose that upon ATP hydrolysis by Ino80^ATPase^ from SHL −6, INO80 rotates around nucleosome by almost 180° to place Ino80^ATPase^ at around SHL −1, so it is poised to translocate DNA from SHL −2. In this model, (i) INO80 relocation away from SHL −6 and DNA translocation from SHL −2 both rely on the ATP-dependent DNA translocase activity of Ino80^ATPase^ and (ii) the two different outcomes from SHL −6 versus SHL −2 arise from differences in how stably the rest of INO80 is anchored at these locations.

## Conclusions and Implications

Our findings that (i) ATP hydrolysis shifts a substantial population of INO80 to a new orientation with Ino80^ATPase^ near SHL −1, (ii) Ino80^ATPase^ bound near SHL-1 could dynamically reach SHL −2 and (iii) a gap placed between SHL −6 and SHL −2 accumulates at SHL −2 with ATP hydrolysis, together provide new evidence in favor of the reorientation model (Figure 1b). We therefore propose that in terms of where nucleosomal DNA is translocated, INO80 behaves like all other remodelers studied to date. Why then does INO80 predominantly engage the nucleosome with Ino80^ATPase^ positioned near SHL −6 before ATP hydrolysis? There is some evidence that other remodelers such as the SRCAP complex, a homolog of *S*.*c*. SWR complex, the Swi2/Snf2 ATPase of the *S*.*c*. SWI/SNF complex and *S*.*c*. Chd1 can analogously engage nucleosomes near the entry-exit site (SHL −8 to SHL −6)^28-31^. However, these cases are substantially different from INO80 because in all these cases, the predominant binding mode without ATP involves the ATPase engaged at SHL −2/+2 (ref 21,23,32).

Prior work provides some possible explanations for these differences. It has been shown that INO80 activity has a unique switch-like sensitivity to flanking DNA that is mediated by the Arp8 module. Recognition of flanking DNA length by the Arp8 module occurs when Ino80^ATPase^ is engaged near SHL −6. ISWI family remodelers also have a flanking DNA length sensing step and recognize flanking DNA via the HAND-SANT-SLIDE domain but do so with their ATPase domains positioned at SHL −2. We therefore propose that INO80’s initial location on the nucleosome serves an additional purpose. Previous work has also shown that unpeeling of DNA near SHL −6 correlates with higher activity on nucleosomal substrates^18^. Such DNA unpeeling exposes the H3 αN helix and allows access to residues such as H3K36, H3T45 and H3K56. Acetylation of H3K36 and H3K56 and phosphorylation of H3T45 are associated with key steps in transcription, DNA replication and DNA repair, all processes where INO80 plays critical regulatory roles^33-38^. We therefore speculate that INO80’s initial location on the nucleosome allows its activity to be responsive to flanking DNA length and nucleosome modifications important for transcription, replication and repair (Figure 5b). The additional ATP dependent reorientation step may then serve as a proof-reading step, ensuring higher specificity of action than what is possible based solely on differences in INO80’s binding affinity for specifically modified nucleosomes^39,40^.

Our work also showcases how the most active states of a remodeler may not always be the most highly populated when visualized by cryoEM in the absence of ATP hydrolysis. Importantly, as described above, uncovering intermediate conformational states for INO80 substantially changed mechanistic models derived from structures obtained in the absence of ATP. The analysis that led to these insights was facilitated by applying cryoROLE to depict the landscape of relative orientation between INO80 and the nucleosome. Methodologically, other currently available methods of analyzing conformational dynamics are unable to tackle such large domain rotations, since these can no longer be considered as small perturbations to a well-defined consensus conformation. We anticipate that similar approaches applied to other chromatin regulators may uncover new steps in their mechanism.

## Methods

### Expression and purification of INO80

The endogenous *S. cerevisiae* INO80 complex was purified as previously described^18,25^. Briefly, a 2×FLAG-tagged INO80 strain (S288C INO80-FLAG) was cultured in YPD medium at 30°C to saturation, and cells were harvested for protein purification. INO80 was first purified by anti-FLAG immunoprecipitation. The elution was then purified by anion exchange chromatography on a Mono Q 5/50 GL column using a linear salt gradient from 100 mM to 1 M KCl over 20 column volumes. Peak fractions were dialyzed into storage buffer containing 25 mM HEPES (pH 7.5), 100 mM KCl, 10% glycerol, mM EDTA, 1 mM DTT, and 0.02% NP-40.

### Reconstitution of gap nucleosomes and nucleosomes with acidic patch mutations

Recombinant *X. laevis* histones were expressed in BL21(DE3) plysS cells and purified as previously described^41,42^. Briefly, DNA containing the Widom 601 nucleosome positioning sequence was amplified by PCR from a plasmid template using a fluorescently labeled primer (IDT)^26^. Large-scale PCR products were separated on a 5% polyacrylamide gel, and the desired DNA band was excised. The gel slice was crushed, soaked overnight in 1× TE buffer, and filtered through a 0.22-μm filter. DNA was ethanol precipitated and dissolved in 1× TE buffer. Fluorescently labeled DNA containing a single nucleotide gap near SHL −2 (25 bp from the dyad toward the flanking DNA), SHL −3 (36 bp), or SHL −6 (56 bp) was generated as previously reported^17^. Refolding of histone octamers was performed as described previously. Nucleosomes were assembled using salt gradient dialysis and purified using a 10 to 30% glycerol gradient.

### ATPase assay

ATPase assays were performed under multiple turnover conditions (nucleosomes and ATP in excess of INO80). Experimentally it was determined that 160 µ M ATP/MgCl_2_ was saturating for all conditions. Reactions were performed with 10 nM INO80 and 100 nM nucleosomes in 26.5 mM Tris pH 7.5, 13.5 mM HEPES pH 7.5, 50 mM KCl, 0.5 mM EDTA, 1 mM MgCl_2_ and trace amounts of γ-^32^P-ATP, at 30°C. Each reaction was incubated at 30°C for 10 minutes and was started with the addition of ATP/MgCl_2_. The no ATP control was taken at the last time point of the reaction. Samples of the reactions were quenched at various timepoints with equal volumes of 50 mM Trips pH 7.5, 3% SDS, and 100 mM EDTA. Inorganic phosphate was separated from ATP on PEI-cellulose TLC plates with 0.5 M LiCl, 1 M formic acid. These plates were exposed overnight on a phosphoscreen and scanned on a Typhoon Imager (GE Life Sciences). The fraction of ATP hydrolyzed was quantified using ImageJ, and initial rates were determined by linear fitting of time points in which less than 10% of ATP was hydrolyzed.

### Preparation of amine-functionalized graphene oxide (GO) grids

Graphene oxide (GO) grids were prepared as previously described^43,44^. Briefly, in a glass petri dish (60 mm in diameter, 15 mm in height) an epoxy coated stainless steel mesh stand was placed at the bottom and DI water was filled to the top. 300 Mesh, R1.2/1.3 Au Quantifoil grids were placed on the mesh stand with carbon side facing upward. Using a syringe, the GO solution (230 μl in total volume) was spread onto the water surface. After draining the water, the GO-coated grids were dried at room temperature for use. GO-covered grids were then submerged in 10 mM ethylenediamine solution diluted in dimethyl sulfoxide (DMSO) and incubated for 5 hours at room temperature. The grids were washed twice with DMSO without ethylenediamine, twice with autoclaved water, twice with ethanol, and dried under ambient conditions. Amino modified grids were stored dry at −20°C until use.

### Electron microscopy sample preparation and data collection

WT INO80 and nucleosomes (Gap −2.5, Gap −3.6, Gap −5.6, or acidic patch mutant (APM)) were mixed at a 2:1 molar ratio and dialyzed into remodeling buffer (26.5 mM Tris pH 7.5, 13.5 mM HEPES pH 7.5, 50 mM KCl, 1.1 mM MgCl_2_, 0.5 mM EDTA, and 2% glycerol) for 2 h. For apo samples, 3 μL of the mixture was directly used for cryo-EM grid preparation following dialysis. The remaining sample was incubated with 1 mM ATP·MgCl_2_ at 30 °C for 30 min before grid preparation. All samples were prepared on amino-functionalized GO grids. Plunge freezing of the grids was carried out by applying 3 μL of sample at 8°C and 100% humidity on FEI Vitrobot IV with a wait time of 4 s and blot force of 0, using ø 55/20 mm blotting filter paper from Ted Pella.

All cryo-EM datasets except the INO80/Gap-3.6 ATP dataset were collected using SerialEM on a Titan Krios G2 equipped with an X-FEG, a Gatan K3 camera, and a Gatan GIF energy filter. The microscope was operated at 300 kV, with an energy filter slit of 20eV. The INO80/Gap-3.6 ATP dataset was collected using SerialEM on a Titan Krios G3i equipped with a C-FEG, a Thermo Scientific Falcon 4i camera, and a Thermo Scientific Selectris X energy filter. The microscope was operated at 300 kV, with an energy filter slit of 20eV. Images were acquired with a nominal defocus range of −0.8 to −1.8 μm.

For the INO80-Gap-2.5 ATP dataset and the INO80-Gap −3.6 apo dataset, 24,761 and 8,472 movies, respectively, were collected at a nominal magnification of 105 K, resulting in a pixel size of 0.835Å. Movies were dose-fractionated into 80 frames, resulting in a total fluence of ~45.8 electrons/Å^2^. For the INO80-Gap-3.6 ATP dataset, 31,641 movies were collected at a nominal magnification of 165 K, resulting in a pixel size of 0.743Å. Movies were dose-fractionated into 1,400 frames with a total exposure of ~60 electrons/Å^2^. For the INO80-Gap-5.6 ATP dataset and the INO80-APM ATP dataset, 9,134 and 12,316 movies, respectively, were collected at a nominal magnification of 105 K, resulting in a pixel size of 0.8189 Å. Movies were dose-fractionated into 80 frames, resulting in a total fluence of ~47.7 electrons/Å^2^. For the INO80-Gap-5.6 apo dataset, 6,431 movies were collected at a nominal magnification of 105 K, resulting in a pixel size of 0.827Å. Movies were dose-fractionated into 60 frames with a total exposure of ~60 electrons/Å^2^. All dose-fractionated movies were motion corrected and dose-weighted on-the-fly using MotionCor2 (ref 45).

### General description of image processing of all datasets

All INO80-Gap nucleosome datasets were initially processed independently using the same workflow. CTF estimation, template picking, particle extraction, 2D classification, and heterogeneous refinement were performed in cryoSPARC^46^. Template picking was carried out using the previously reported INO80-nucleosome reconstruction (EMDB-28613) as the reference. Particles were extracted using a box size of 448 × 448 pixels for all datasets except the INO80-Gap −3.6 ATP dataset, for which a box size of 480 × 480 pixels was used because of its smaller pixel size. Initial heterogeneous refinement produced well-resolved INO80 density but weak nucleosome density, indicating conformational heterogeneity.

To minimize classification bias, particles from all datasets were merged and imported into RELION^47^ for 3D refinement. A mask containing the nucleosome and Arp5 module was generated to subtract densities outside the mask. Signal-subtracted particles were rescaled from 448 × 448 to 240 × 240 pixels or from 480 × 480 to 280 × 280 pixels prior to 3D classification. Two major classes were identified. For each class, focused refinement of the nucleosome yielded reconstructions at 3.3 Å (Class 1) and 3.0 Å (Class 2). The subtracted particles were then reverted to the original particles, and focused refinement of INO80 resulted in reconstructions at 3.1 Å and 3.0 Å, respectively. Composite maps were generated by aligning the individually refined nucleosome and INO80 reconstructions using the overlapping Arp5 module the common reference. In these final composite reconstructions, Class 1 corresponds to the previously reported conformation with Ino80^ATPase^ bound at SHL −6, whereas Class 2 represents a distinct conformation with Ino80^ATPase^ placed near SHL −1. Particles were subsequently traced back to their original datasets for validation and further analyses.

### Identifying the gap in Gap-3.6 nucleosome

To visualize the location of gap in the dataset of INO80/Gap-3.6 nucleosome sample with ATP, we further processed Class 2 particles, in which Ino80^ATPase^ is located at near SHL −1. A mask containing nucleosomes was generated to subtract signals outside the mask. The particles after signal subtraction were subjected to 3D classification, resulting in a major class with well-resolved histone density and extended exit-side DNA. This class was refined in cisTEM to a resolution of ~3.0 Å and used for atomic model building. Density corresponding to a single nucleotide rather than a base pair of nucleotides at SHL −2 is seen in this structure, clearly indicating a gap.

### Landscape of relative orientation between INO80 and nucleosome

Analysis of relative orientation between INO80 and bound nucleosome were performed using cryoROLE^27^. Particles belonging to Class 1 and Class from all datasets were combined for analysis. During the image processing workflow described above, focused refinement of the INO80 complex was performed using the same INO80 reference orientation for both classes. In contrast, focused refinements of the nucleosome in Class 1 and 2 were performed using two nucleosome reference oriented differently, nearly opposite to one another. Consequently, the two particle sets could not be directly merged for individual refinement of the INO80 complex and nucleosome. Although the merged particles yielded a well-resolved reconstruction of the INO80 complex, individual refinement of the nucleosome did not produce a high-resolution reconstruction because the initial nucleosome reference orientations differed substantially between the two classes.

To overcome this limitation, the rotational transformation between the two nucleosome reference frames was determined by aligning the Class 2 nucleosome to that of Class 1. This transformation was then applied to the Euler angles of all Class 2 nucleosome particles, thereby placing both particle sets into a single common nucleosome reference. The transformed Class 2 particle STAR file was subsequently merged with the Class 1 particle STAR file, and focused refinements of the INO80 complex and nucleosome were performed. As a result, the Euler angles of the INO80 complex and nucleosome were determined with respect to a common INO80 reference and a common nucleosome reference, respectively, and a common composite map can be generated, in which Ino80^ATPase^ is positioned at SHL 0, dyad, which is referred as a = 0° in the landscape. thereby enabling generation of the final combined relative orientation landscape. Particle box sizes used for cryoROLE refinement were identical to those used during the image-processing workflow described above. Scaled Local Density (SLD) is used to estimate local particle population in the landscape, and SLD =1 is used as a threshold to remove particles that are too sparsely distributed in the landscape.

### Image processing of the INO80-APM nucleosome dataset

Template picking was carried out using the previously reported INO80-nucleosome reconstruction (EMDB-28613) as the reference. 527,749 particles were selected after initial cryoSPARC-based processing and then were imported into RELION for 3D refinement. A mask containing the nucleosome and Arp5 module was generated for signal subtraction, and densities outside the mask were subtracted. Signal-subtracted particles were then subjected to focused 3D classification. Two major classes were identified. For Class 1, focused refinement of the nucleosome yielded reconstruction at 6.5Å. The subtracted particles from this class were reverted to the original particles and focused refinement of INO80 resulted in reconstruction at 3.3 Å. For Class 2, the subtracted particles were reverted to the original particles and refinement of the INO80-nucleosome complex resulted in reconstruction at 7Å.

### Model building

For the model building, the initial model was generated by fitting the available coordinates into our cryo-EM density maps by using Chimera. These coordinates include the INO80 core, and the model of the *X*.*l* nucleosome (PDB: 9C9Z and 1KX5). The inconsistent parts were then manually built and refined in coot. The structures were refined using Phenix with secondary structure constraints.

## Data availability

Composite maps and fitted atomic model of INO80-nucleosome with Ino80^ATPase^ at SHL −6 and at SHL −1 are deposited to EMDB and PDB (SHL −6: EMD-78186 and PDB-37HD, SHL −1: EMD-78179 and PDB-37GR).

## Acknowledgement

This work is supported by grants from National Institute of Health (1R35GM140847 to Y.C., R35 GM127020 to G.J.N.). Equipment at the UCSF cryo-EM facility was partially supported by National Institutes of Health (NIH) grants (S10OD020054, S10OD021741, and S10OD025881). Operation of the facility is supported by G. Gilbert and D. Bulkley. Y.C. is an Investigator of Howard Hughes Medical Institute.

## Author Contributions

H.W., U.K., E.N.M., G.J.N. and Y.C. conceptualized the study. E.N.M. generated the DNA gap-containing nucleosomes and developed and implemented the reaction conditions for cryo-EM. E.N.M. and U.K. purified INO80 and performed biochemical assays with nucleosomes. H.W. carried out most of the cryo-EM related studies of INO80/Gap and INO80/APM complexes, with substantial assistance from U.K. for cryo-EM studies of the INO80/APM complex. C.L. and Y.C. conceptualized landscape analysis. H.W. and C.L. carried out landscape analysis and optimized cryoROLE procedure. H.W., U.K., G.J.N. and Y.C. wrote the manuscript, with input and feedback from all authors.

## Correspondence and requests for materials

should be addressed to Geeta J. Narlikar and Yifan Cheng.

---

**Supplementary Figure 1.**
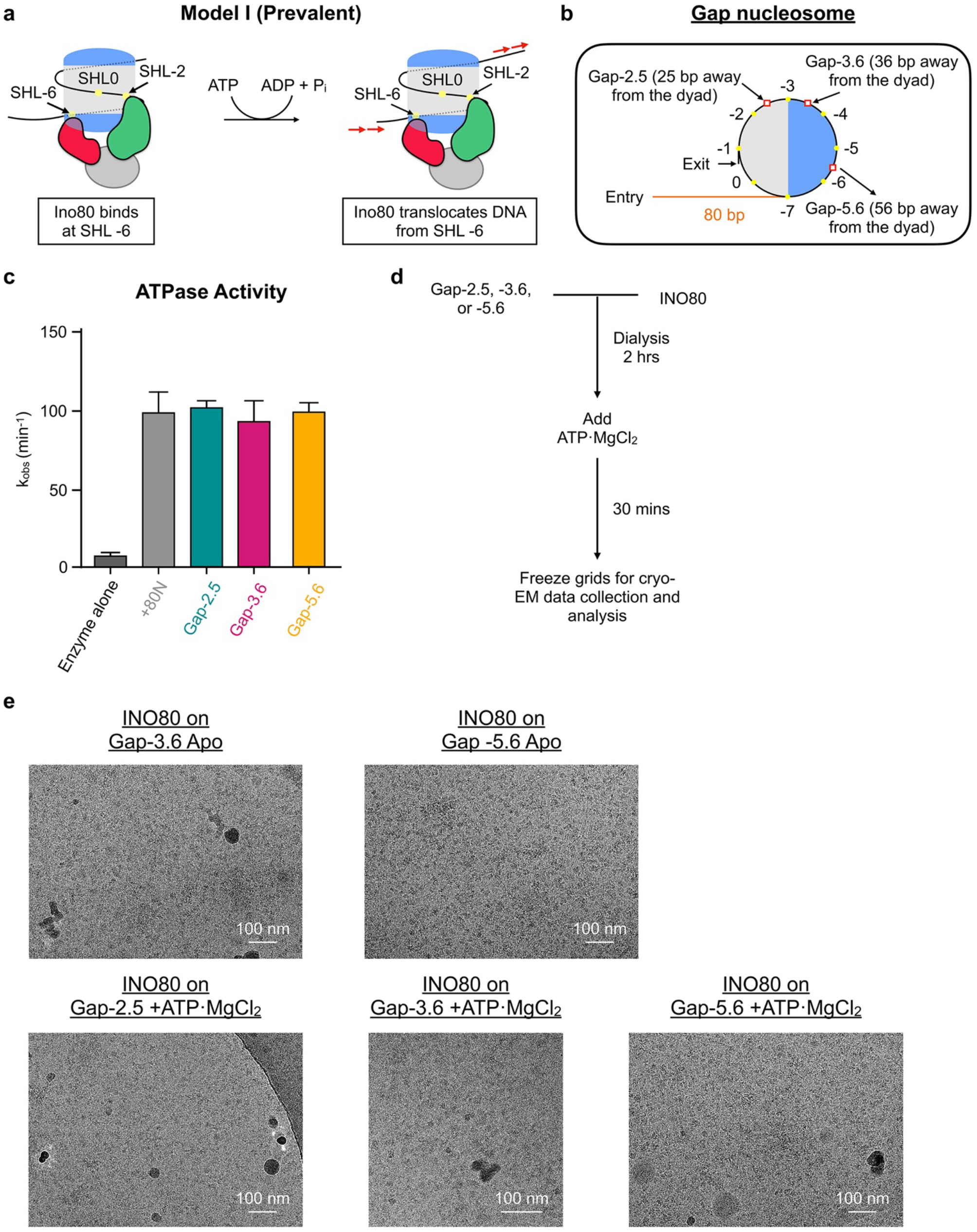
Preparation of INO80/Gap nucleosome samples. **a)** Previous model of INO80-mediated nucleosome remodeling. INO80 binds nucleosomes at SHL −6 and directly translocates DNA from this position. **b)** Schematic of a +80 nucleosome containing DNA gaps at SHL −2.5, −3.6, or −5.6. Gap positions are labeled by red boxes. **c)** ATPase activity of INO80 on WT nucleosome, Gap-2.5, Gap-3.6, and Gap −5.6 nucleosomes. **d)** Workflow for preparation of cryo-EM samples. **e**, Representative cryo-EM micrographs of the INO80-Gap nucleosome samples.

**Supplementary Figure 2.**
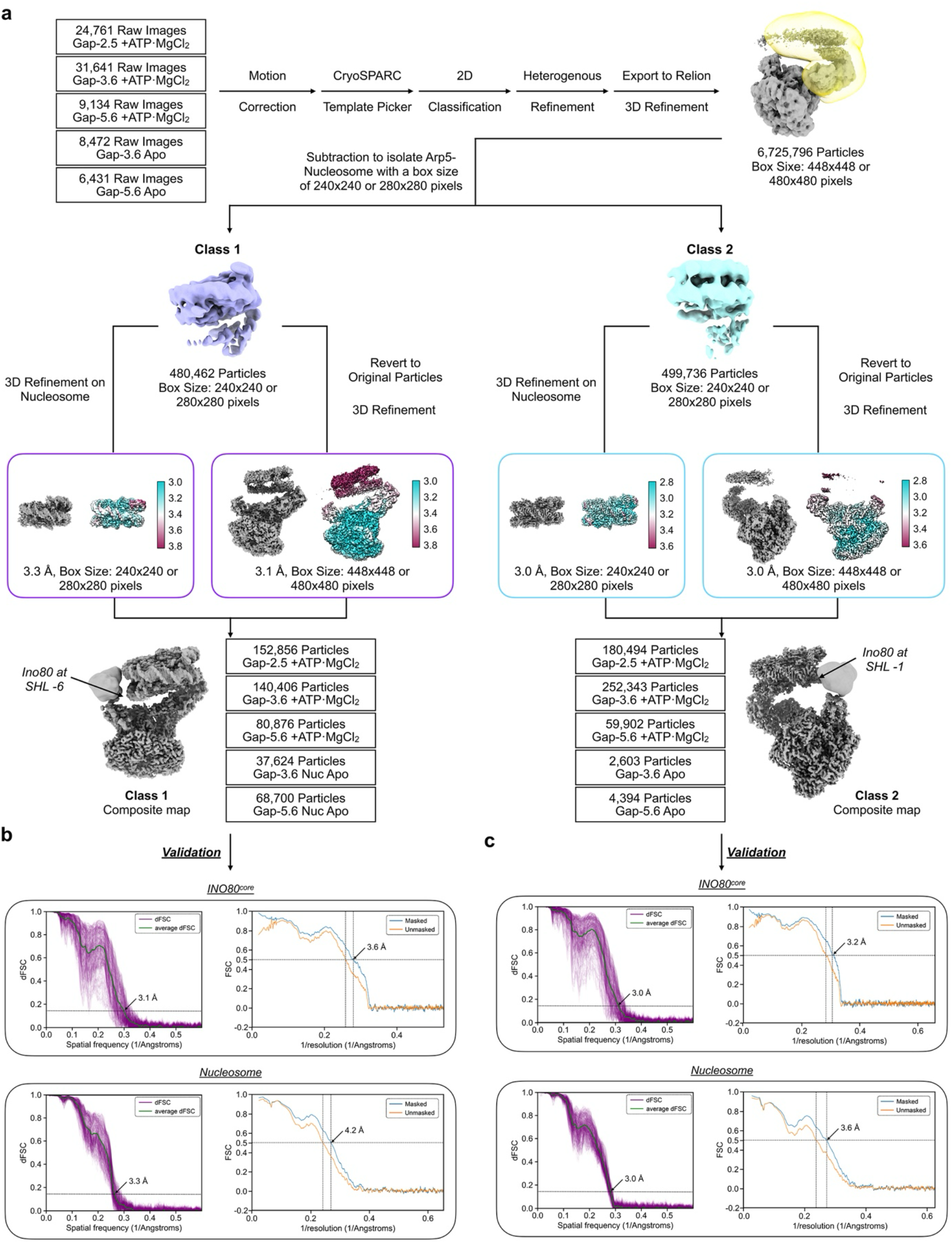
Cryo-EM image processing of the INO80/gapped nucleosome datasets. **a)** Image processing workflow for the combined INO80/gap nucleosome cryo-EM datasets. Composite maps are generated following the same procedure. **b)** Resolution estimation from directional Fourier shell correlation (dFSC, 0.143 criteron, left) and model-to-map FSC curves (FSC = 0.5 criterion) of INO80 core (upper) and nucleosome (bottom) of Class 1 composite map. **c)** Resolution estimation from dFSC (left) and model-to-map FSC (right) of INO80 core (upper) and nucleosome (bottom) of Class 2 composite map. Estimated resolutions are marked.

**Supplementary Figure 3.**
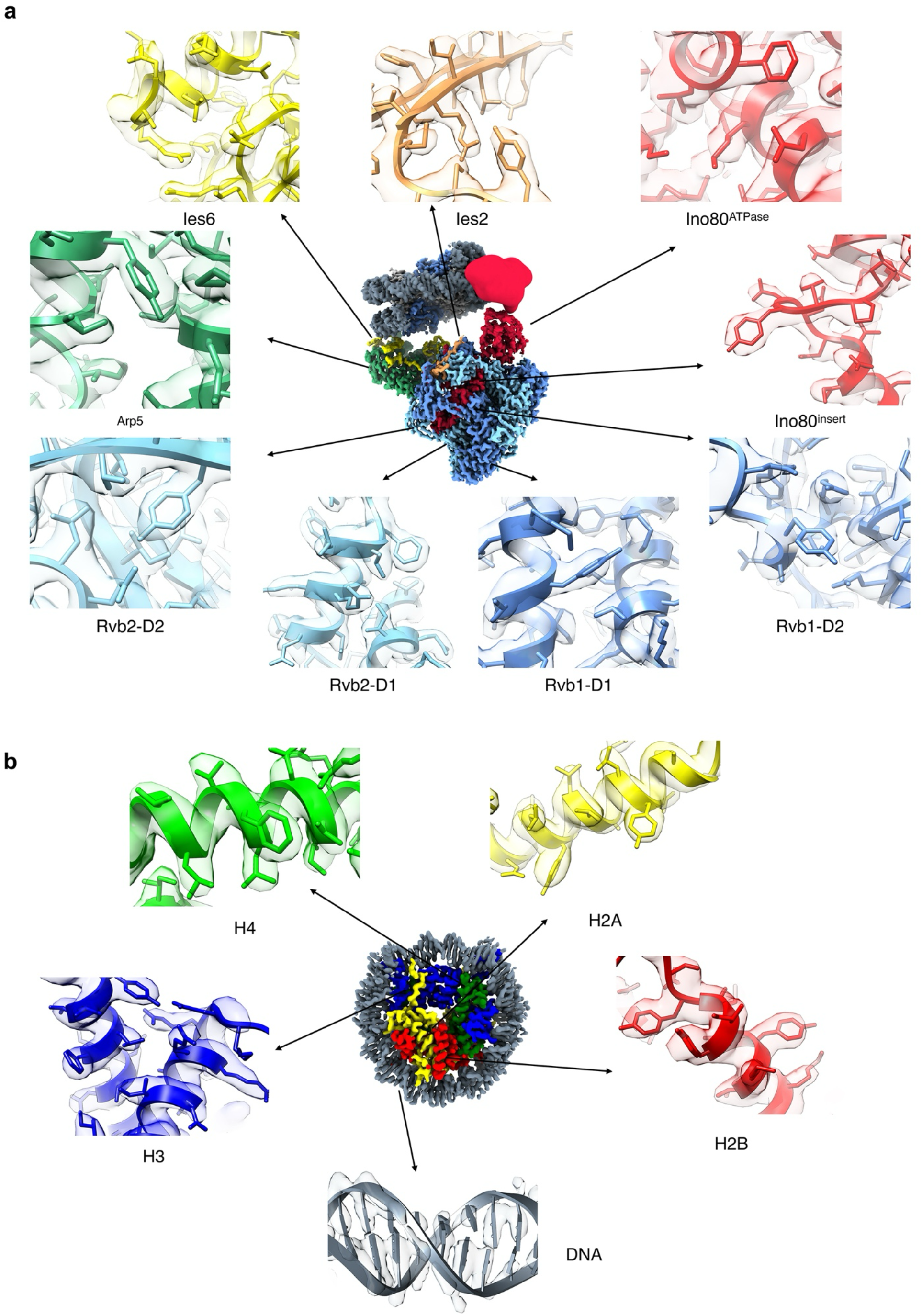
Representative densities of the INO80-Gap nucleosome complex. Representative densities of Class 2 composite map. **a**) INO80 core and **b)** nucleosome.

**Supplementary Figure 4.**
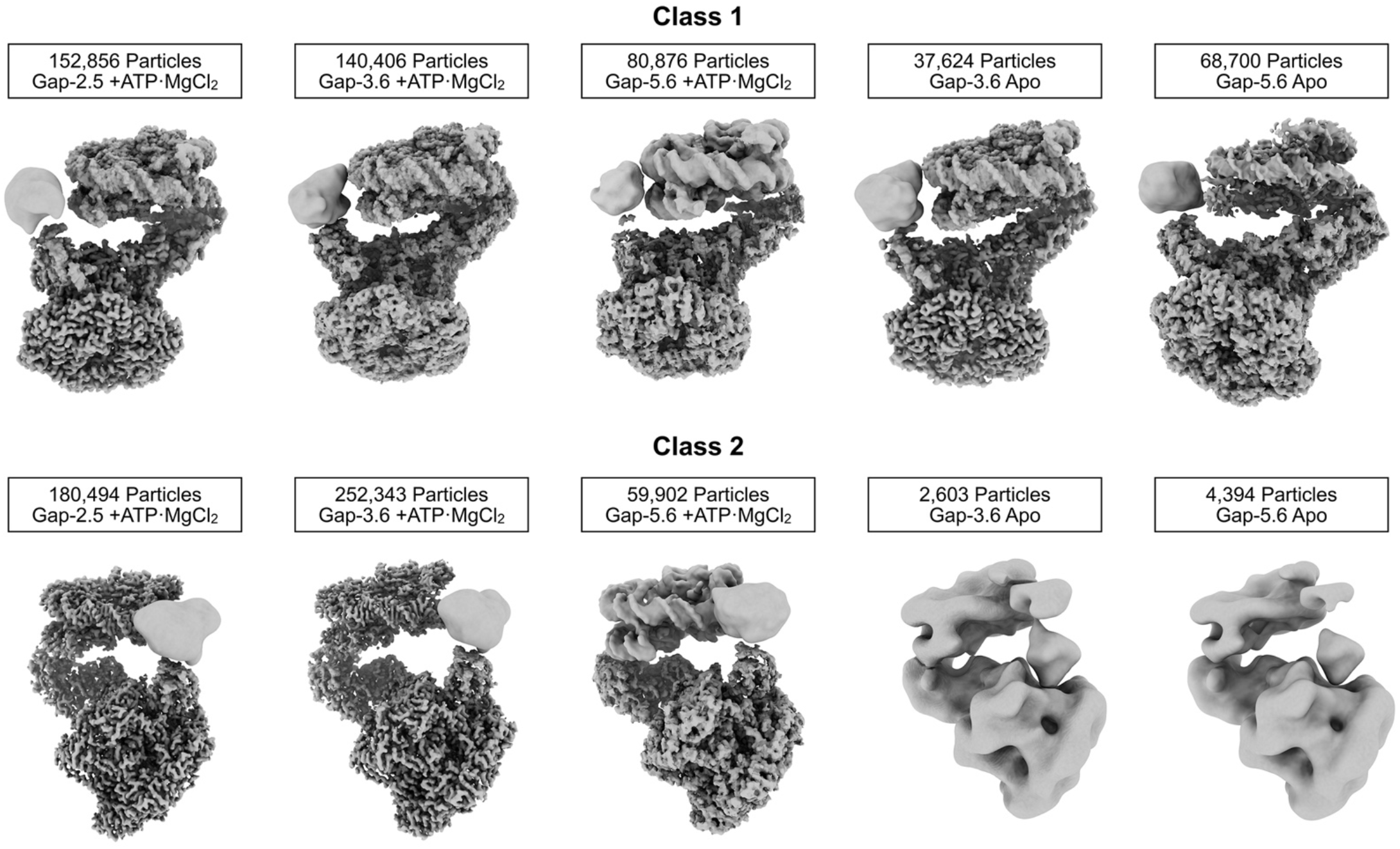
Individual composite maps of Class 1 and 2 from each dataset. Composite maps of Class 1 (upper row) and Class 2 (bottom row) of individual datasets, as marked. Particle numbers for each reconstruction are marked. All maps are displayed in the same orientation.

**Supplementary Figure 5.**
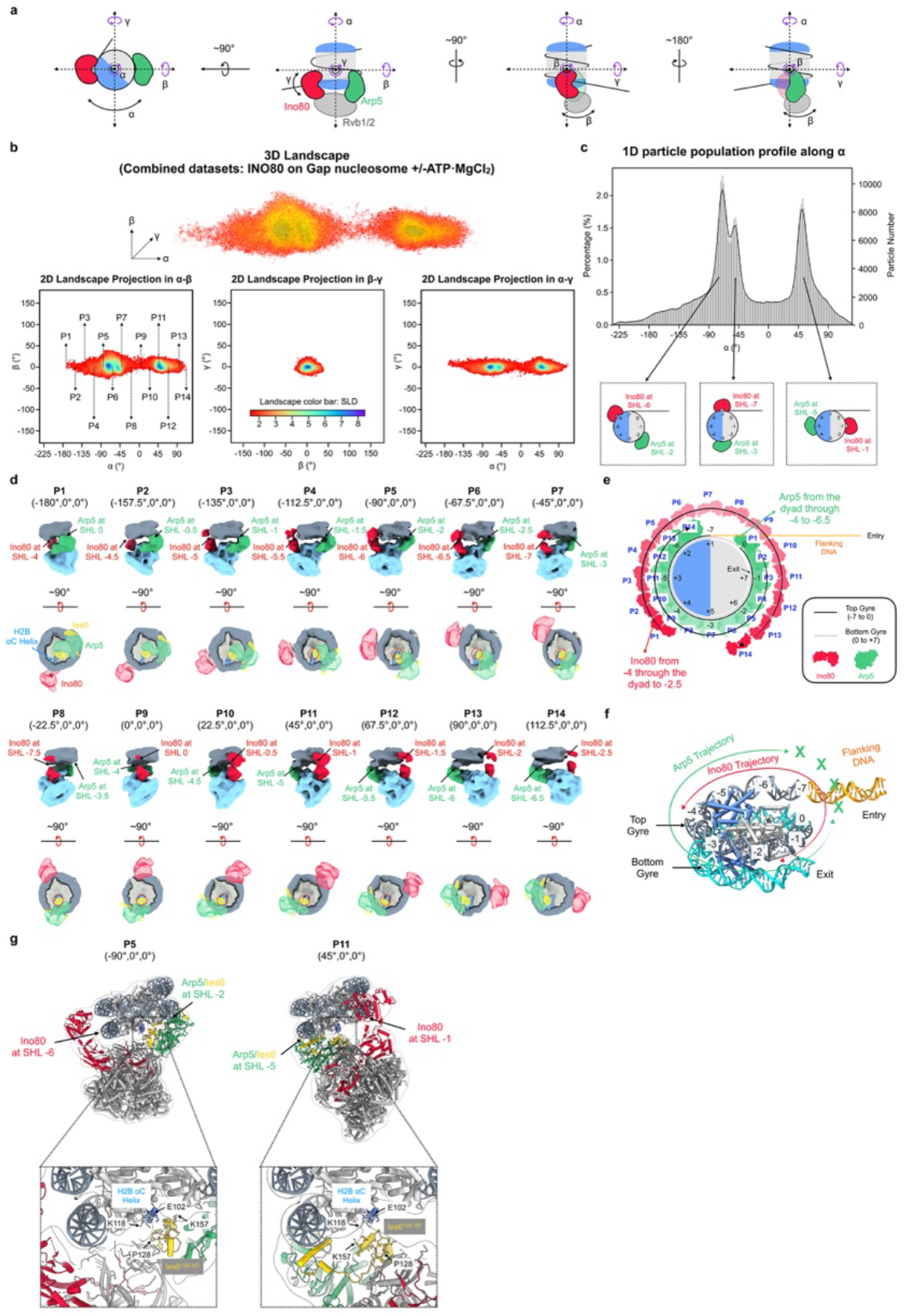
The relative orientation landscape of all datasets combined, including Gap-2.5 ATP, Gap-3.6 ATP, Gap-5.6 ATP, Gap-3.6 Apo, Gap-5.6 Apo. **a)** Schematic illustration of α, β, and γ angles in the canonicalized coordinate frame used to define the relative orientation between INO80 core and nucleosome. **b)** Upper panel: A 3D view of the landscape, with the direction of rotational axes marked. Each dot in the landscape represents a particle and its coordinate represent the relative orientation between INO80 core and nucleosome of that specific particle, and color coded with scaled local density (SLD). Particles with SLD less than 1 are filtered out. Bottom panel: 2D projection views of the 3D landscape onto the α-β, β-γ, and α-γ planes are shown. SLD = 1.5 was used to display projections. 14 positions with 22.5° intervals along a axis are marked in the α-β projection. **c)** Particle distribution profile along α axis derived from the landscape. Peaks are labeled as the location (SHL) of Ino80^ATPase^ (red) and Arp5 (green) on nucleosome. **d)** Reconstructions calculated from particle extracted from the landscape at position marked as P1 to P14 along the α axis. The upper panels show side views, and the lower panels show top views from the direction of INO80 core, in which only densities of part of Ino80^ATPase^ (red), Arp5 (sea green) and Ies6 (yellow) are shown. Densities of histone are in light grey DNA in dark grey. H2B αC helix of acidic patch is marked in cornflower blue. Red dotted circle marks the existence of a density connecting Ies6 and the H2B αC helix. **e)** Trajectories of Ino80^ATPase^ (red) and Arp5 (green) illustrated on nucleosome with SHL position marked. The positions of Ino80^ATPase^ and Arp5 are derived from reconstructions shown in (**d**). **f**) An illustration shows that the moving trajectory Ino80^ATPase^ (red) and Arp5 (green) while keeping Arp5 on top gyre of nucleosomal DNA. **g)** Structural comparison of representative reconstructions corresponding to P5 (left, Ino80^ATPase^ at SHL −6) and P11 (right, Ino80^ATPase^ at SHL −1). Densities are displayed in low contour level as transparent. Insets highlight the interaction between the H2B αC helix and Ies2.

**Supplementary Figure 6.**
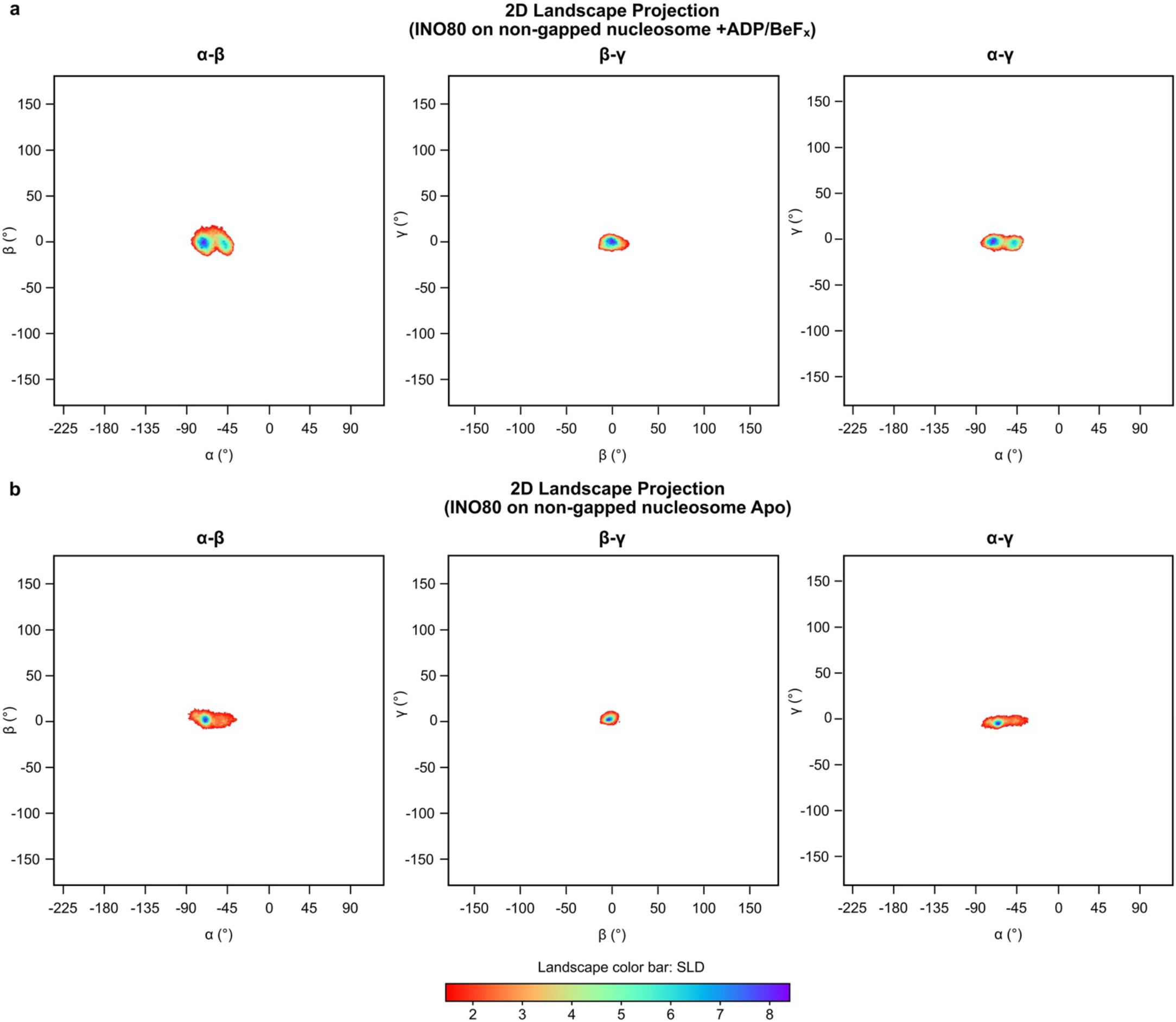
Landscapes of the INO80/non-gapped nucleosome. **a)** 2D projections of the landscape of INO80/non-gapped nucleosome in the presence of ADP/BeF_x_ onto the α-β, β-γ, and α-γ planes. **b)** 2D projections of the landscape of INO80/non-gapped nucleosome in the apo state onto the α-β, β-γ, and α-γ planes.

**Supplementary Figure 7.**
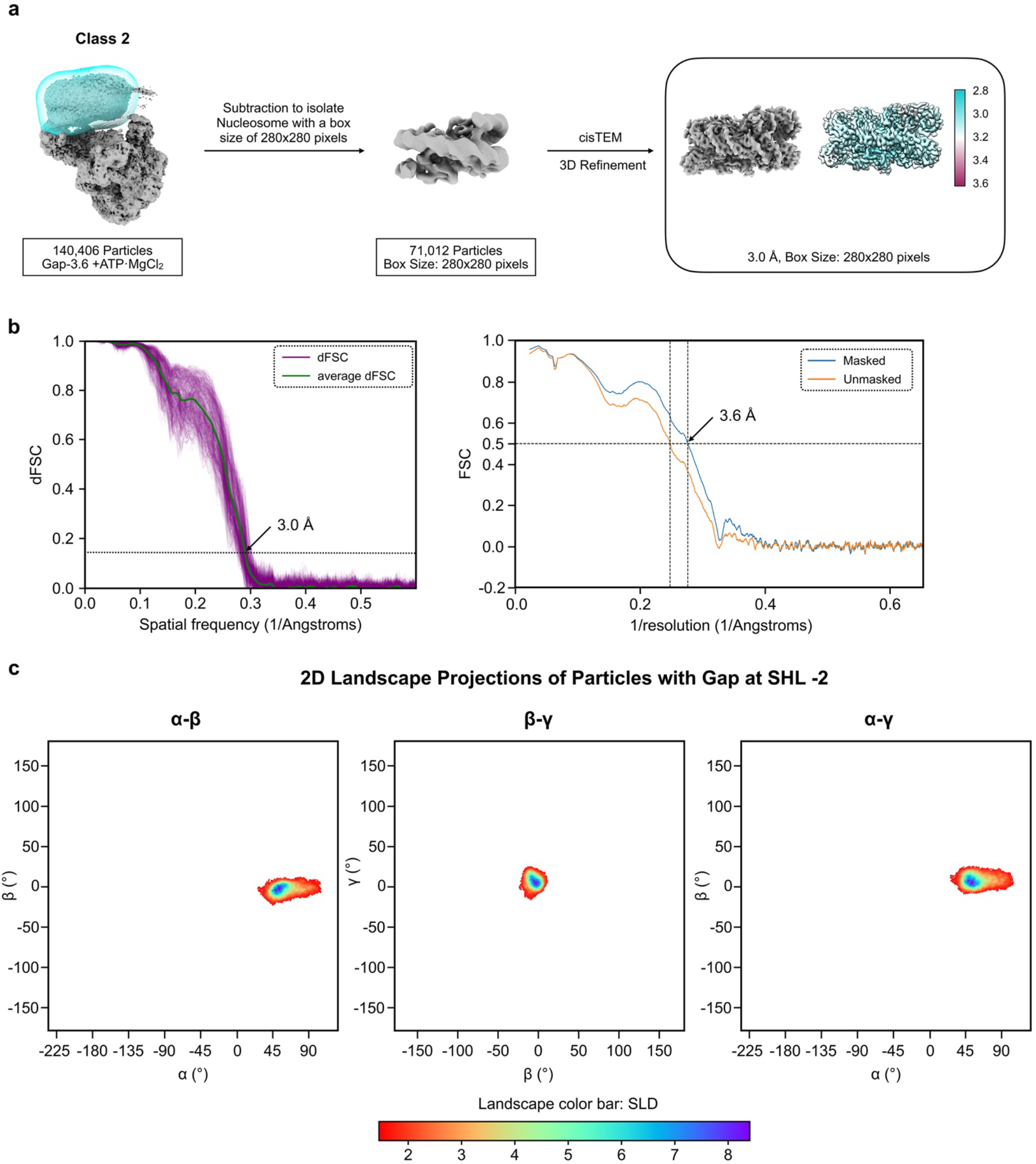
Image Processing and landscape of particles with gap at SHL −2. **a)** Image processing workflow of the Class 2 particle subset of the INO80/Gap-3.6 ATP dataset to identify particles with gap at SHL −2. **b)** Resolution estimation of the refined nucleosome map. dFSC and model to map FSC curves are shown. **c)** 2D projections in α-β, β-γ, and α-γ planes of the landscape of particles with gap at SHL −2.

**Supplementary Figure 8.**
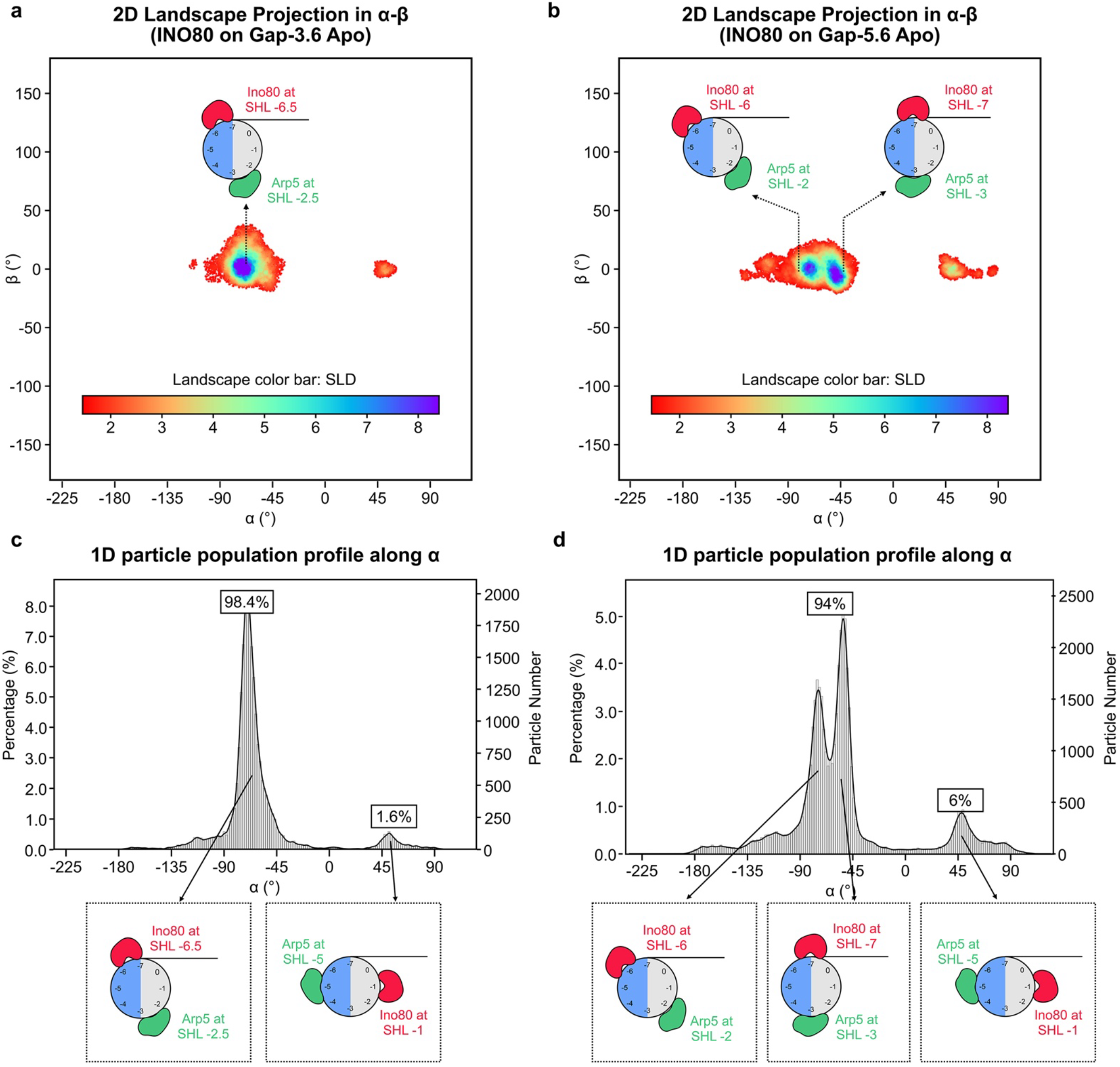
Landscape of INO80/Gap-3.6 and INO80/Gap-5.6 in apo state. **a)** and **b)** 2D projections of 3D landscapes onto the α-β plane for the INO80/Gap-3.6 apo (**a**) and INO80/Gap-5.6 apo (**b**). Orientations between INO80 core and nucleosome are illustrated as cartoon. Landscapes are colored by SLD as defined by the color bar. **c)** and **d)** Particle distribution profile along α derived from the landscapes shown in **(a)** and **(b)**. Percentage of particles under the peaks are marked. Insets show the ATPase and Arp5 binding positions corresponding to the peaks in the profile.

**Supplementary Figure 9.**
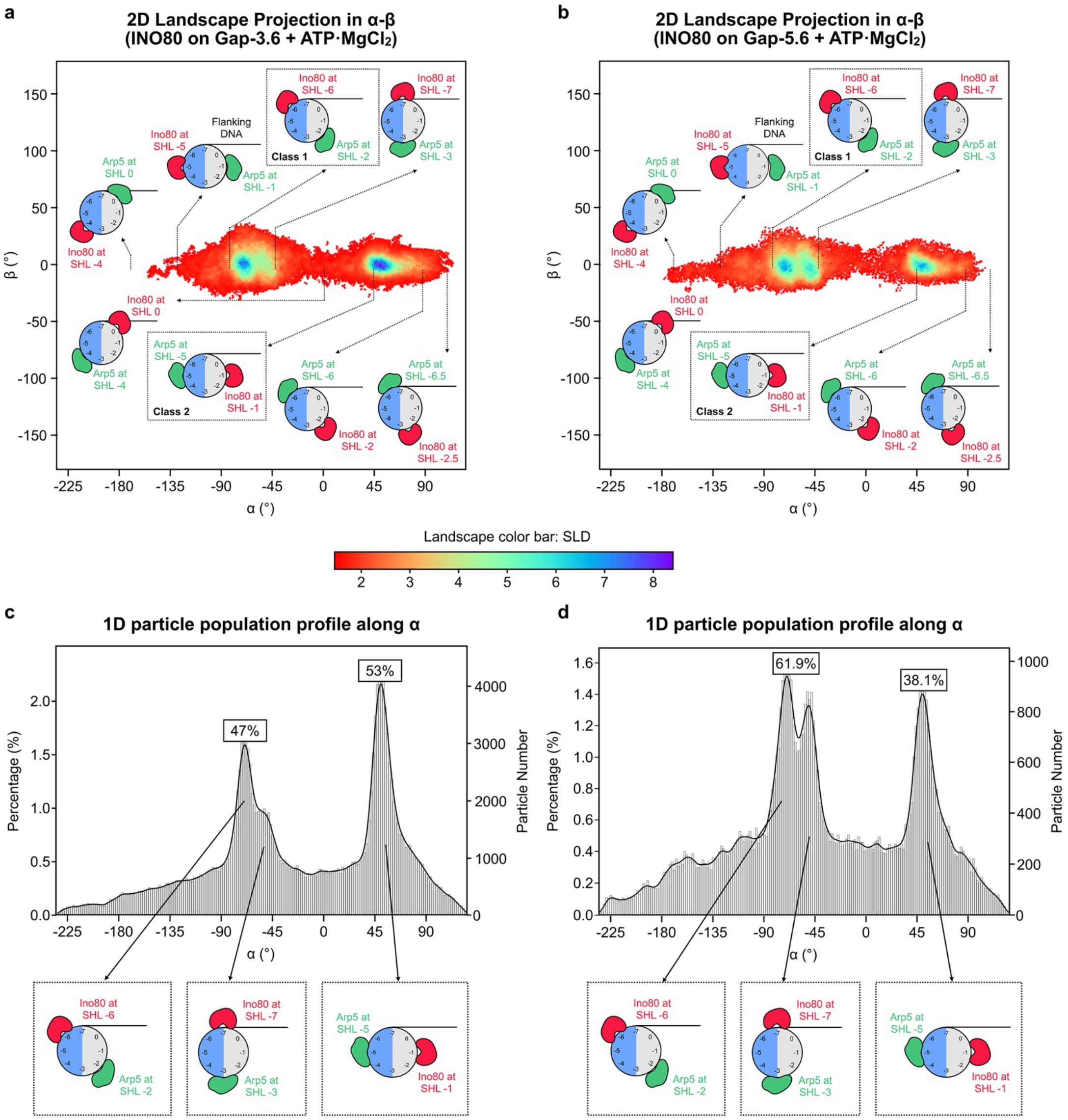
Landscape of INO80/Gap-3.6 and INO80/Gap-5.6 in the presence of ATP. **a)** and **b)** 2D projections of 3D landscapes onto the α-β plane for the INO80/Gap-3.6 with ATP (**a**) and INO80/Gap-5.6 with ATP (**b**). Orientations between INO80 core and nucleosome from selected locations in the landscape are illustrated as cartoon. Landscapes are colored by SLD as defined by the color bar. **c)** and **d)** Particle distribution profile along α derived from the landscapes shown in **(a)** and **(b)**. Insets show the ATPase and Arp5 binding positions corresponding to the peaks in the profiles.

**Supplementary Figure 10.**
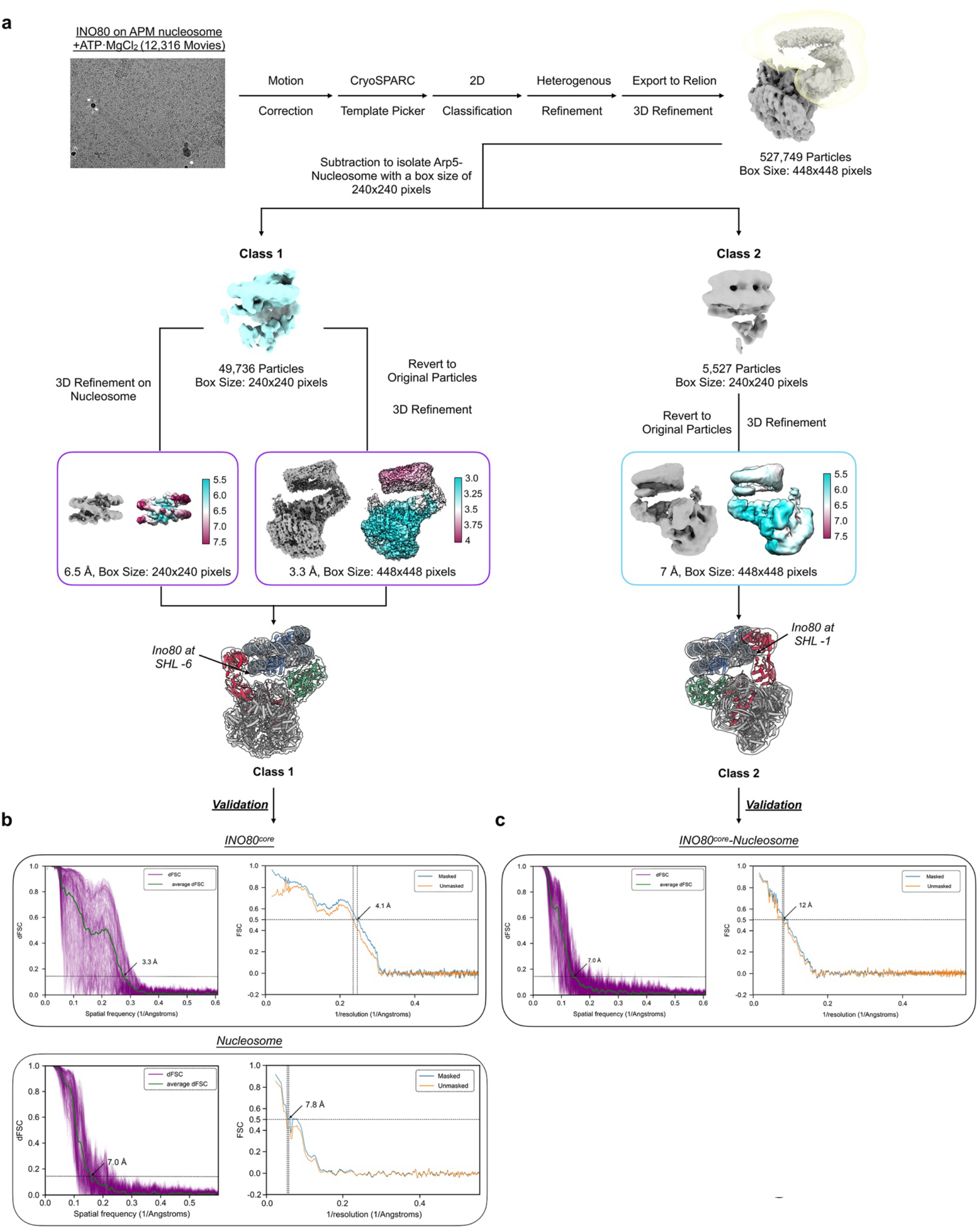
Image processing flowchart of INO80/APM nucleosome with ATP. **a)** Image processing flowchart of INO80/APM nucleosome with ATP. **b**) Resolution estimation of Class 1 INO80 and nucleosome from dFSC (left, 0.143 criterion) and model-to-map FSC (right, 0.5 criterion). **c**) dFSC (left, 0.143 criterion) and model-to-map FSC (right, 0.5 criterion) of Class 2 reconstruction.

**Supplementary Figure 11.**
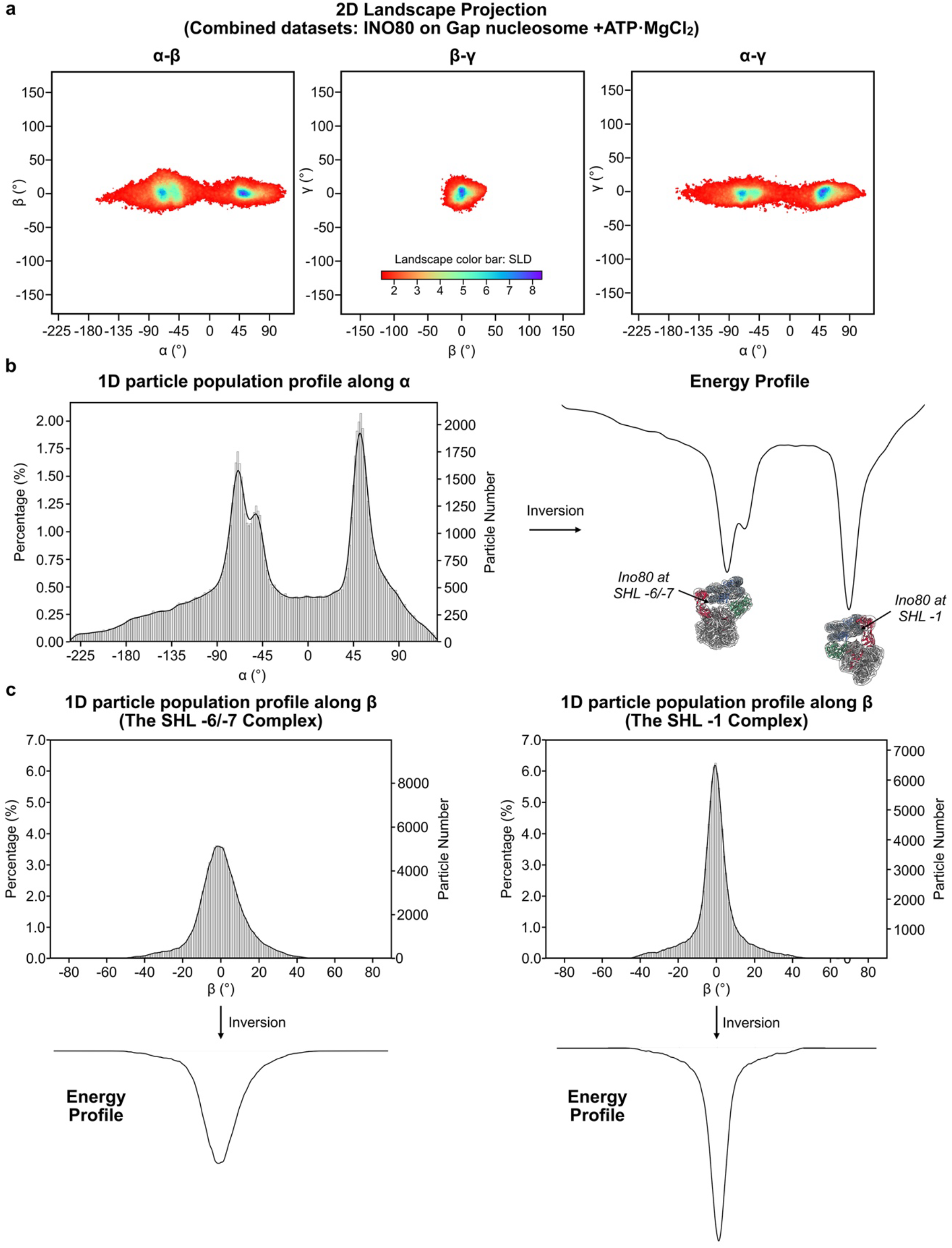
Free energy landscapes predicted from the relative orientation landscape. **a)** 2D projections of the relative orientation landscape of the combined INO80/Gap nucleosome ATP datasets onto the α-β, β-γ, and α-γ planes. **b)** Particle distribution profile along α (left) and a conformational free energy landscape generated assuming that the population distribution represents a near equilibrium distribution (right). With this assumption, particle distribution at various conformation is estimated to be inversely correlated with the free energy landscape and inversion of the particle distribution profile produces a corresponding conformational free energy landscape. **c)** Particle distribution profile along β axis for particles with Ino80^ATPase^ at near SHL −6/−7 complex (left) and the SHL −1 complex (right). Together, they suggest that INO80 has a more stable interaction with nucleosome when Ino80^ATPase^ is at near SHL −1 than that at SHL −6/−7.

**Supplementary movie 1: 3D view of relative orientation landscape**

Movie shows 3D landscape with all particles included, transition to a landscape views with a SLD threshold of 1 and 1.5.

**Supplementary movie 2: 3D view of landscape with SLD threshold 1.5**

Movie shows 3D landscape with a SLD threshold set to 1.5, followed by reconstructions calculated from particles extracted from the landscape.

